# Inversely regulated immune-related processes mediate anxiety-obesity links in zebrafish

**DOI:** 10.1101/2023.02.08.527622

**Authors:** Hila Yehuda, Nimrod Madrer, Doron Goldberg, Hermona Soreq, Ari Meerson

## Abstract

Anxiety disorders often associate with metabolic impairments, but the underlying developmental and molecular mechanisms are yet unknown. To seek RNAs that may link anxiety and obesity, we subjected RNA from zebrafish larvae of a caffeine-induced anxiety model and a high fat diet (HFD)-induced obesity model to RNA-sequencing. We found differentially expressed genes in the larval anxiety and obesity models, including long noncoding RNAs and transfer fragment RNAs. Surprisingly, they were inversely regulated and comprised overrepresentation of immune system pathways, e.g., interleukin signaling and inflammation. Similarly, inverse regulation persisted in adulthood, but with different overrepresented immune system processes, e.g., T cell activation, leukocyte cell-cell adhesion and antigen processing and presentation. Furthermore, unlike the known link in adult zebrafish, obesity in zebrafish larvae was not accompanied by anxiety-like behavior. These results may reflect an antagonistic pleiotropic phenomenon involving re-adjusted modulation of the anxiety-metabolic links with the immune system. Furthermore, the HFD potential to normalize the anxiety-upregulated immune-related genes may explain previously reported protective roles of high fat diet in rodent anxiety and Alzheimer’s disease models.

## Introduction

Anxiety is an adaptive response for preventing future harm (Robinson *et al*, 2019). However, in excess and when persistent, anxiety reduces the quality of life and is considered to be a psychiatric condition, belonging to the heterogenous group termed anxiety disorders (Robinson *et al*, 2019). Anxiety disorders are the largest group of mental disorders in the Western world (Craske *et al*, 2017), with a 20% lifetime occurrence (Meier & Deckert, 2019). The COVID pandemic has been accompanied by yet a greater number of anxiety cases (Walsh *et al*, 2021), highlighting the need to explore the origins of infection-induced anxiety and seek ways to treat these devastating disorders and the Long COVID symptoms (Sudre *et al*, 2021). Intriguingly, anxiety disorders have often been found to be positively linked to obesity and the metabolic syndrome (characterized by three of the following five symptoms - type II diabetes, hyperlipidemia, high LDL, low HDL and hypertension (Huang, 2009)); this is the case in humans (Van Reedt Dortland *et al*, 2013; Tang *et al*, 2017), rats (Dutheil *et al*, 2016), mice (Xia *et al*, 2021; Almeida-Suhett *et al*, 2017; Meydan *et al*, 2016) and adult zebrafish (Türkoğlu *et al*, 2022; Picolo *et al*, 2021). Furthermore, anxiety disorders present 30-50% heritability, although specific genes or loci have yet to be found (Shimada-Sugimoto et al., 2015), and obesity is causally related to 17 human obesity loci-related genes, assessed as such in *C. elegans* (Ke *et al*, 2021). The shared links between anxiety and obesity/metabolic syndrome (Baker *et al*, 2017; Nousen *et al*, 2014; Hiles *et al*, 2016) would predictably imply that genes hyperactivated in one disorder may be further elevated in the other.

Although social factors may be partially responsible for the link between obesity and anxiety, such factors cannot explain the high incidence of anxiety detected among obese persons (Baker *et al*, 2017). Moreover, animal studies show definite associations between anxiety disorders and obesity/metabolic syndrome. Obese high fat diet (HFD)-fed rats with higher fasting glucose levels than matched controls demonstrate anxiety-like symptoms (Dutheil *et al*, 2016). Similarly, overweight HFD-fed mice display anxiety-like behavior (Xia *et al*, 2021; Gainey *et al*, 2016; Ogrodnik *et al*, 2019), although in one study the correlation between obesity and anxiety disappeared when attaining a certain weight (Ogrodnik *et al*, 2019). Correspondingly, selectively bred obesity-prone Sprague-Dawley rats had greater anxiety-like behavior than obesity-resistant ones, but only after weight gain, regardless of their diet (Alonso-Caraballo *et al*, 2019). Also, HFD did not lead to anxiety-like behavior in obesity-resistant rats, even though the diet brought about weight gain and increased adiposity that was comparable to the weight gain found normally in obesity-prone rats (Alonso-Caraballo *et al*, 2019); this indicates that genetics or other factors also play a part in obesity-induced anxiety. Notably, while HFD-fed adult zebrafish showed anxiety-like behavior relative to controls fed a regular diet (Türkoğlu *et al*, 2022), their five-day old HFD-fed offspring did not exhibit anxiety-like behavior (Türkoğlu *et al*, 2022), hinting that obesity may correlate with anxiety-like behavior in vertebrates in an age-related manner.

Zebrafish have become a model for many diseases, since they are highly homologous to mammals, including humans, in many respects (Nguyen *et al*, 2013; Lieschke & Currie, 2007). Genetically, 70% of the human genome has an orthologue in the genome of zebrafish, and 82% of known human morbid genes are related to at least one zebrafish orthologue (Howe *et al*, 2013). Zebrafish gastrointestinal (Zang *et al*, 2018; López Nadal *et al*, 2020), neural (Fontana *et al*, 2018), endocrine (Zang *et al*, 2018) and immune systems (Novoa & Figueras, 2012; de Abreu *et al*, 2018) have similar mechanisms of action to those of mammals. Likewise, lipid storage and insulin regulation are conserved, with obese zebrafish demonstrating a similar dysregulation of lipid metabolism to that in humans (Zang *et al*, 2018). The anatomy, gene expression and projections between brain regions of the teleost brain is similar to that of other vertebrates (Friedrich *et al*, 2010). As in humans, and unlike rodents, the zebrafish stress hormone is cortisol (Fontana *et al*, 2018). Its increase is associated with anxiety in both zebrafish (Egan *et al*, 2009) and humans (Lenze *et al*, 2011; Blay & Marinho, 2012). The immune system of zebrafish has features similar to mammals, with many of the same cytokines, chemokines and immune cell types (Novoa & Figueras, 2012). Thus, the zebrafish is a good model to examine links between anxiety and metabolic disorders.

One manner of identifying links between anxiety and obesity would be to examine differentially expressed (DE) genes in both disorders. Such genes may be protein coding genes, which make up less than 2% of the human genome (Aprile *et al*, 2020) and/or genes transcribed to noncoding RNAs, including microRNAs (miRNAs) and long noncoding RNAs (lncRNAs), which can take part in regulation of anxiety (Issler & Chen, 2015; Kolshus *et al*, 2014; Murphy & Singewald, 2019; Meydan *et al*, 2016; Cui *et al*, 2017; Spadaro *et al*, 2015) and metabolic disorders (Iacomino & Siani, 2017; Arner & Kulyté, 2015; Aryal *et al*, 2017; Meydan *et al*, 2016; Lo *et al*, 2018; Cai *et al*, 2018). Recently, a small noncoding RNA family, transfer RNA fragments (tRFs) has acquired new recognition (Zorbaz *et al*, 2022). As the name insinuates, these fragments are derived from enzymatic fragmentation of transfer RNAs, which occurs on pre-tRNAs and mature tRNAs. This enzymatic cleavage is not random. Under stress, the anticodon is cleaved by angiogenin in mammals or by other enzymes in other species, resulting in 5’ and 3’ tRNA halves (also termed tiRNAs) (Mathew *et al*, 2022). Under other circumstances (e.g. cancer), RNase Z cleaves the pre-tRNAs, producing tRF-1s, and the dicer and other enzymes (some still unknown) divide the mature tRNAs into many types of tRFs, i.e., 5’ tRFs (tRF-5), 3’ tRFs (tRF-3) and internal tRFs (i-tRFs) (Kim *et al*, 2020; Mathew *et al*, 2022; Rosace *et al*, 2020). tRFs, especially the 5’ halves, have been found to take part in immune system reactions (Zorbaz *et al*, 2022; Winek *et al*, 2020) and have roles in neurological disorders (Mathew *et al*, 2022).

To date and to the best of our knowledge, no single RNA-seq study sought coding or noncoding RNAs (ncRNA) that link metabolic disorders with anxiety disorders. Therefore, the main goal of this research was to challenge the concept that specific RNAs may link anxiety and obesity and identify related DE transcripts.

## Results

### Experimental validation of the anxiety and obesity models

To investigate transcriptomic profile modifications induced in anxiety and obesity, 15-day post-fertilization (dpf) zebrafish larvae were used in two well-known models. The anxiety model was based on a short exposure (0.5 h) to a high caffeine dose (100 mg/liter) (Richendrfer *et al*, 2012; Rosa *et al*, 2018; Tran *et al*, 2017), and the obesity model was induced by an egg yolk-based high fat diet (Zang *et al*, 2018). Caffeine-exposed larvae (CAF), compared to non-treated larvae (NO), showed greater anxiety-like behavior, including increased thigmotaxis (duration in center - means were 7.4% CAF vs 24.9% NO, Welch Two Sample t-Test, p=0.002) and erratic swimming (cumulative average absolute turn angle [degrees/min] were 31.2 for CAF vs 21.2 for NO, Welch Two Sample t-Test, p=0.00008) (Supplementary Figures S1A, S1B, S1E), as well as higher cortisol levels (means were CAF – 120, NO – 55 pg/larvae, Welch Two Sample t-Test, p=0.002) (Figure S1F). High cortisol levels are known to correlate with anxiety in zebrafish (Egan *et al*, 2009). The caffeine-exposed larvae also had reduced locomotion (Figure S1C), in agreement with published studies (Tran *et al*, 2017; Rosa *et al*, 2018). In the obesity model, two independent experiments (n=24-25) revealed a greater percentage of adipocyte-laden larvae among high fat diet (HFD)-fed larvae compared to standard diet (SD)-fed larvae (56% vs 21%, respectively). In addition, HFD-fed larvae had more abdominal adipocytes than did SD-fed larvae (2.12 vs 0.50 adipocytes/larva, respectively; Welch Two Sample t-Test, p=0.006) (Figure S1G), and the average total area of adipocytes in HFD-fed larvae was larger than in SD-fed larvae (1973 µm^2^ versus 222 µm^2^, respectively; Welch Two Sample t-Test, p=0.0008) (Figure S1H). Together, these results validate the efficacy of the anxiety and obesity models.

### Most transcripts upregulated in anxious larvae were downregulated in obese larvae

The concept of RNAs linking anxiety and obesity and identifying those DE genes was explored by subjecting RNA from larvae of the two models to next generation RNA-sequencing. The RNA sequence of each model comprised control and treatment sets: for the anxiety model - NO versus CAF, and for the obesity model - SD versus HFD, with 30 larvae per replicate and three replicates per treatment.

Following RSEM annotation and expression calculation, DESeq2 was used for normalization and statistics. According to DESeq2, among the 32520 poly(A)+ genes, including all mRNA and many lncRNA genes, there were 29720 genes with a non-zero read count. Principal component analysis (PCA) showed that the levels of all (32520) poly(A)+ transcripts clustered according to treatment (Figure 1A, B). Thus, the biological replicates had similar expression profiles, and these profiles differed among treatments. The two controls, SD and NO, probably varied due to larvae being transferred to wells containing filters for quick removal following caffeine exposure, which was not carried out on the SD (and HFD) larvae.

**Figure 1.**
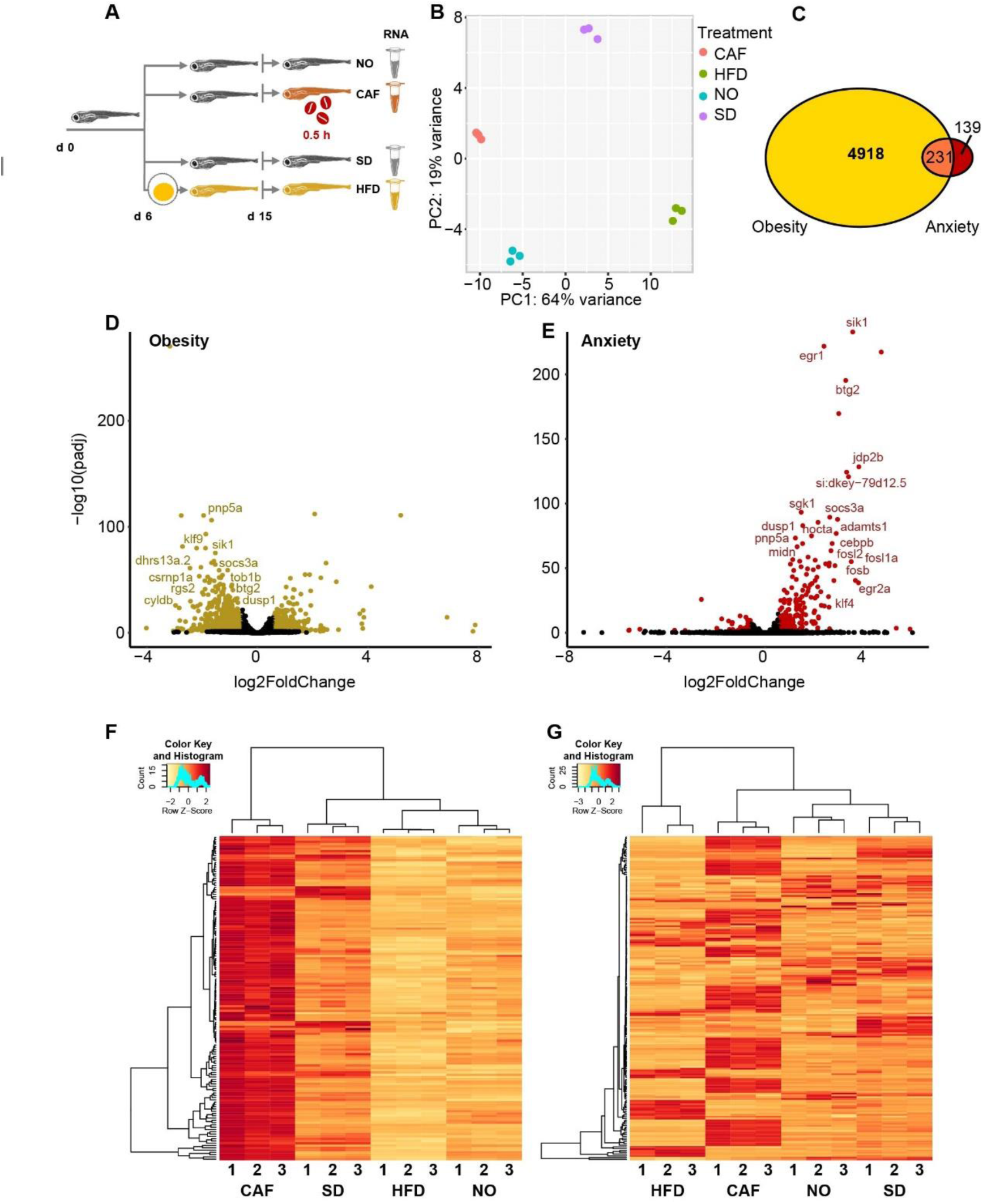
Most anxiety-upregulated larvae transcripts were downregulated in obesity. A Experimental design-zebrafish larvae (6 dpf) were fed either standard diet (SD) or high fat diet (HFD, based on hard-boiled egg yolk solution). At age 15 dpf, a group of the SD-fed larvae were further divided into caffeine-exposed larvae (CAF) and their controls (NO). Following caffeine exposure (0.5 h) of the CAF larvae, larvae of all treatments were snap frozen until RNA extraction for RNA-seq. B Profiles based on 32520 poly (A)+ mapped transcripts clustered according to treatment in the PCA (B). C A total of 231 poly(A)+ transcripts were DE (padj<0.05) in both the obesity (yellow) and anxiety (red) models. D Obesity model volcano plot - DE, transcripts color-marked gold; padj<0.05 and absolute log2 fold change>0.58 [>1.5 and <0.67-fold change]). Labeled transcripts are among the 107 genes upregulated >1.5 fold in the anxiety model and downregulated <0.67 fold in the obesity model. E Anxiety model volcano plot – DE, transcripts color-marked red; padj<0.05 and absolute log2 fold change>0.58 [>1.5 and <0.67-fold change]). Labeled transcripts are among the 107 genes upregulated >1.5 fold in the anxiety model and downregulated <0.67 fold in the obesity model. F Heatmap of transcripts per million (TPM) from RSEM of 142 DE poly(A)+ transcripts upregulated in anxiety model (>1.5 fold, padj<0.05) G Heatmap of 182 DE isoform transcripts (FDR<0.05) that were upregulated >1.5-fold in the anxiety model and downregulated <0.67-fold in the obesity model (30 larvae per replicate, three replicates per treatment)

There were 5149 (17.3%) DE genes in the obesity model compared to merely 370 (1.2%) in the anxiety model (padj<0.05)(Figure 1C). The difference likely reflects the longer duration of the obesity model induction (10 days) versus that of the anxiety model (0.5 h). Additionally, this large difference in the fraction of DE genes in the two models may indicate a greater effect of obesity versus anxiety on gene expression. Furthermore, DE genes in the obesity model were mostly downregulated, whereas DE genes in the anxiety model were mostly upregulated (Figures 1D, 1E). This inverse regulation was even more prominent when analyzing the 231 genes commonly DE (padj<0.05) in both models (Figure 1C), i.e., their direction of change was almost totally inverse: 142 genes were upregulated in the anxiety model, but only 10 genes were upregulated in the obesity model (fold change > 1.5). Inversely, 143 genes were downregulated in the obesity model, with only 23 genes being downregulated in the anxiety model (fold change <0.67). Thus, among the 142 genes that were upregulated in the anxiety model, 107 (75%) were downregulated in the obesity model. These DESeq2-analyzed results are reflected in the heatmap (Figure 1F). Of the 231 intersecting genes, only one gene was upregulated in both models (foxq1a, fold change > 1.5), and 8 genes were downregulated in both (si:ch211-125o16.4, pglyrp6, osgn1, apoda.2, ins, urgcp, arid6 and CABZ01046997.1, fold change < 0.67). We conclude that fish larvae show inverse profiles of DE genes in the obesity and anxiety states.

### DE isoform results were similar to DE gene results

DESeq2 analyzes the expression of genes, but not that of isoforms. If one isoform of a gene is upregulated and another isoform of the same gene is downregulated to the same degree, the DE information is lost, since the two changes cancel each other out. According to EBSeq analysis, there were 52061 mapped isoform transcripts, including mRNAs (41203), retained intron long noncoding (lncRNAs (3107)), processed transcript lncRNAs (2841), long intergenic noncoding RNAs (lincRNAs (1901)), antisense lncRNAs (754), sense intronic lncRNAs (54), sense-overlapping (9) and other ncRNAs (Table S2), and of these mapped isoform transcripts, 10688 were DE (FDR<0.05). Among the DE transcripts, 473 were DE isoform transcripts that belonged to the selected DE patterns, appropriate for DE in the anxiety and obesity models (Materials and Methods). Of these, 182 had a fold change of either > 1.5 or < 0.67 in both models, clustering according to treatment (Figure 1G). Similar to the gene transcriptomics analysis above, 92 of the 182 isoforms (51%) that were upregulated in the caffeine-exposed larvae were downregulated in the HFD-fed larvae. Thus, isoform transcripts, as well, showed inverse regulation patterns in anxious and obese fish larvae.

### Inflammation/immune system protein-coding transcripts are upregulated in anxious larvae and downregulated in obese larvae

Next, we explored the overrepresented gene ontology (GO) terms (Fisher’s Exact Test, FDR<0.05) of the anxiety-upregulated [238 mapped/281] and downregulated [73/89] transcripts, and the obesity-upregulated [1993/2396] and downregulated [2331/2753] transcripts, padj<0.05) ones. Most notable among the anxiety-upregulated and obesity-downregulated terms were the immune-related protein class, “basic leucine zipper transcription factor” (involved in the development of immune cells (Yin *et al*, 2017)) and the three immune system-related pathways: “inflammation mediated by chemokine and cytokine signaling pathway”, “CCKR (gastrin- and cholecystokinin-mediated regulation of cellular processes) signaling map” and “interleukin signaling pathway” (Figure 2A); The latter two were also overrepresented when the fold change threshold of the genes included in the lists was changed to >1.5 fold and <0.67 fold. In comparison, digestion associated Reactome Pathways, i.e., “metabolism of water-soluble vitamins and cofactors”, “degradation of the extracellular matrix” and “digestion” were common among the two downregulated gene lists (Figure S2A). Comparing the two upregulated lists revealed that the highest fold-change GO-slim biological process (BP) terms comprised “small molecule biosynthetic process” and “response to organonitrogen compound” (Figure S2B). The only commonly overrepresented term in the anxiety-downregulated and obesity-upregulated comparison was the GO-Slim BP “macromolecule metabolic process”(Figure S2C). More results derived from the comparison of the anxiety-upregulated and obesity-downregulated genes included the GO-slim BP terms, “response to peptide hormone”, “cellular response to hormone stimulus” and “alcohol biosynthetic process”. We conclude that the dominant change in both experimental models involved changes in immune/inflammatory regulation, which were exacerbated under anxiety and suppressed in obesity.

**Figure 2.**
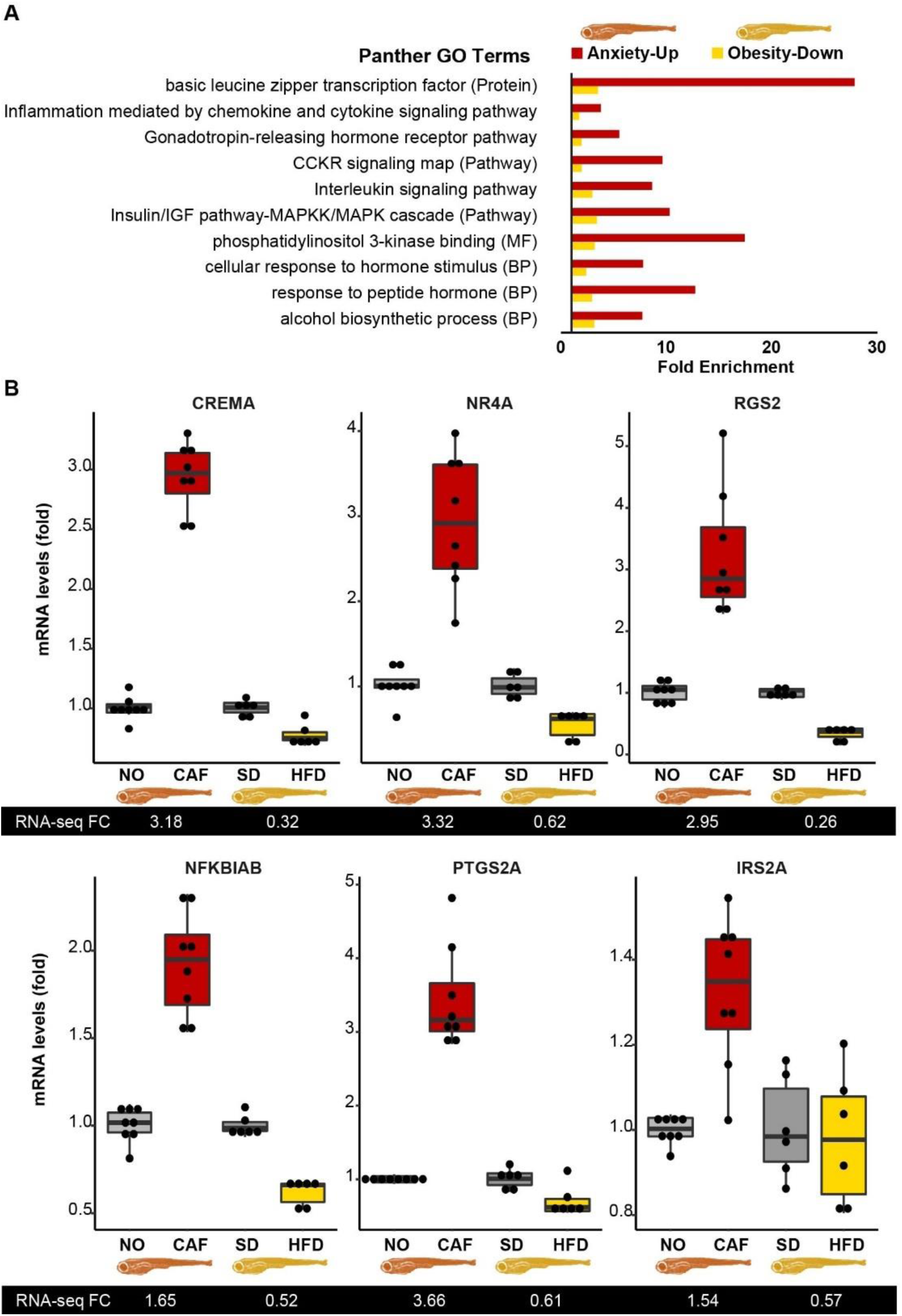
Larval inflammation/immune pathways are upregulated in anxiety and downregulated in obesity. A PANTHER Overrepresentation Test identified “CCKR - gastrin- and cholecystokinin-mediated regulation of cellular processes signaling map”, “inflammation mediated by chemokine and cytokine signaling pathway” and “interleukin signaling pathway”. Reference genes were all those included in the Danio rerio database, Fisher’s exact test, correction by FDR<0.05. B qRT-PCR results of the anxiety and obesity models are presented as graphs, with RNA-seq results below each graph, n= 6-8 replicates (10-30 larvae/replicate), 2-3 separate experiments, Student’s t-Test. Reference genes are the average of eef1a1l1 and actb2. p<0.02 - for all models except the obesity model of IRS2A.

The immune/inflammatory-related pathways noted above included a total of 99 genes, with 14 (13%) being among the inversely regulated 107 anxiety-obesity genes, i.e. early growth response 1 (egr1, also a cholinergic transcription factor in innate immune-related cells (Madrer & Soreq, 2020)), nuclear receptor subfamily 4 group A member 1 (nr4a1 aka NUR77), serpin-family E member 1 (serpine1), cAMP responsive element modulator a (crema), regulator of G protein signaling (rgs)13, prostaglandin-endoperoxide synthase 2a (ptgs2a aka COX2a), rgs2, NFκB inhibitor α b (nfkbiab), insulin receptor substrate (irs)2a, irs2b, chemokine [C-C motif] receptor 9a (ccr9a), MYC proto-oncogene (mycb), interleukin 4 receptor, tandem duplicate 1 (il4r.1) and chemokine [C-X-C motif] receptor 4b (cxcr4b), in at least one of the pathways. Six of the 14 inflammation/immune-related DE transcripts were tested and successfully validated by qRT-PCR (Figure 2B). Most of these transcripts may suppress and/or induce inflammation, depending on the conditions (Welty *et al*, 2016; Francés *et al*, 2015; Rodríguez-Calvo *et al*, 2017; Catrysse & van Loo, 2017; Cai & Liu, 2012; McNabb *et al*, 2020; Zhu *et al*, 2020). The seemingly inverse roles of these regulators may hence be subject to additional control (e. g., via non-coding RNAs) under stress conditions.

The remaining 93 genes were enriched for biological processes related to ectoderm, mesoderm and endoderm formation (83-fold), negative regulation of MAPK cascade (27-fold), positive regulation of proteolysis (18-fold), regulation of protein serine/threonine kinase activity (16-fold), MAPK cascade (8-fold), regulation of cell cycle (5-fold) and regulation of transcription by RNA polymerase II (3-fold) (Figure S3A) and the p53 Pathway (22-fold) (Figure S3B).

### lincRNAs upregulated in the anxiety model were downregulated in the obesity model

In addition to coding sequences, the 182 DE isoform transcripts in both models (FDR<0.05, fold change >1.5 and <0.67) included 29 noncoding poly(A)+ RNAs, that could be divided into three classes (according to Biomart Ensembl): 12 long intergenic noncoding RNAs (lincRNAs) (41%), 9 processed transcripts (31%) and 8 retained introns (28%) (Figure 3A). Similar to the pattern found for DE poly(A)+ RNAs, most of the lincRNAs that were upregulated in the anxiety model were downregulated in the obesity model. The DE lincRNAs, identified by RNA-seq, were tested by qRT-PCR (Figure 3B), validating FO681323.1-202, AL954191.1-201 and CR926130.2-201 upregulation in the anxiety model and downregulation in the obesity model. CABZ01048956-203, FO834828-201 and FO904966-201 also tended to be inversely regulated. As these molecules exhibited a regulation pattern similar to most DE protein coding genes in this study, and since lincRNAs are known to have regulatory activities (Deniz & Erman, 2017), these molecules may also take part in the regulation of inflammatory pathways. To our knowledge, this is the first report on possible functions of these transcripts.

**Figure 3.**
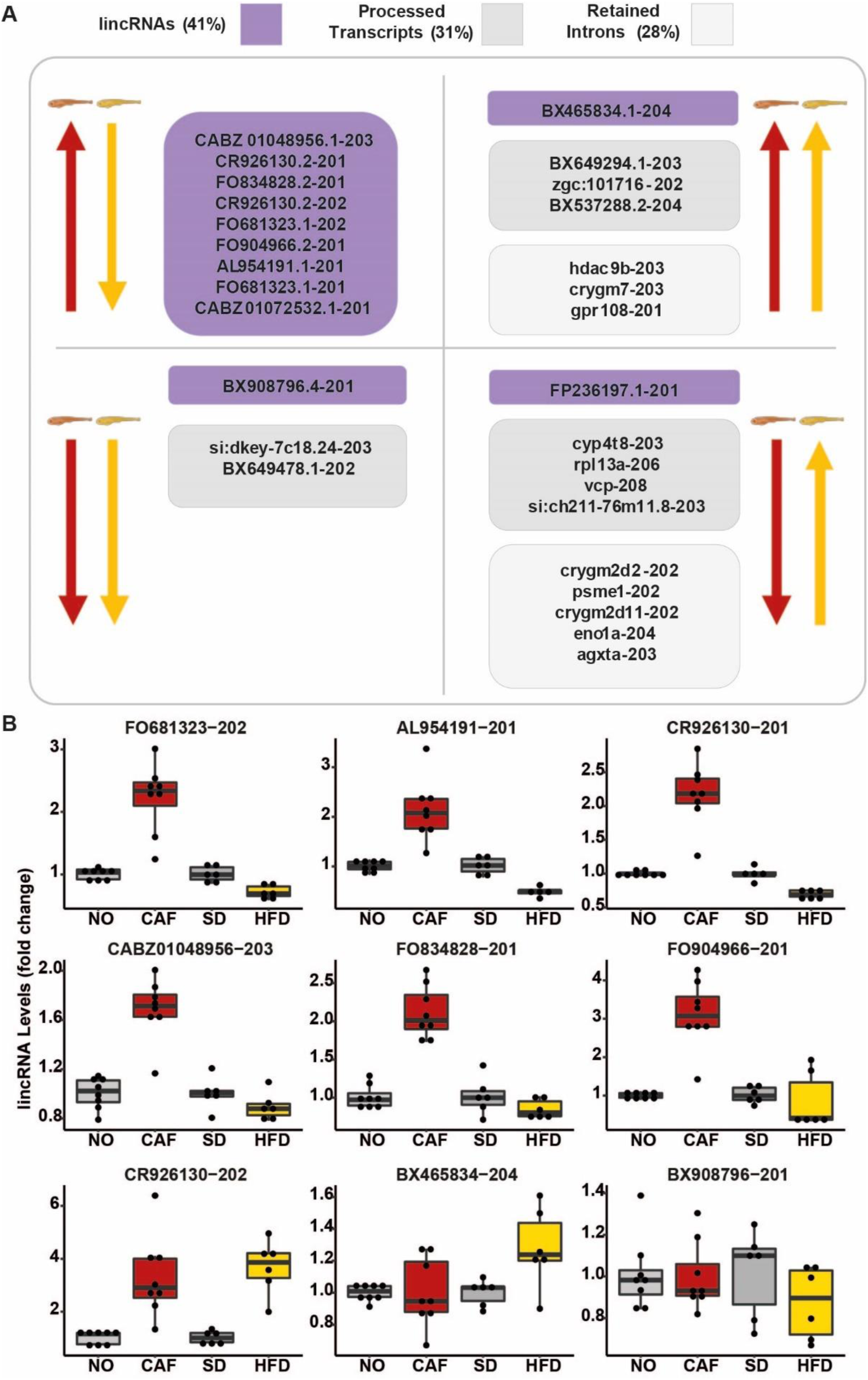
Most DE lincRNAs were upregulated in the anxiety model and downregulated in the obesity model, similar to protein-coding genes. A lncRNAs, classified as lincRNA (purple), processed transcripts (dark grey) and retained introns (light grey) with fold change >1.5 or <0.66, FDR<0.05. Arrows pointing upward: upregulation, arrows pointing downward: downregulation. Yellow: obesity model; red: anxiety model. B qRT-PCR of lincRNA. n= 5-8 replicates (10-30 larvae/replicate), 2-3 separate experiments. Reference genes: average of eef1a1l1 and actb2. p<0.005 for all models for FO681323-202, AL954191-201, CR926130-201 and CR926130-202 and for the anxiety model CABZ01048956-203, FO834828-201 and FO904966-201, Student’s t-Test.

### The processed transcript, si:dkey-7c18.24-203, was downregulated in both the anxiety and the obesity models according to RNA-seq and qRT-PCR

Of the retained intron and processed transcript lncRNAs that were DE according to RNA-seq, only those upregulated in both models or downregulated in both models were tested by qRT-PCR. None of the RNA-seq results of the retained intron lncRNAs (Figures S4A-4B) were confirmed by qRT-PCR. However, one of the processed transcripts (Figures S4C-4F), si:dkey-7c18.24-203, was downregulated by about 30% in both models according to qRT-PCR (Figure S4F), validating its RNA-seq results.

### Neighboring mRNAs of DE lincRNA did not imply functions for lincRNA

Notably, lncRNAs may regulate neighboring mRNAs transcribed from genomic regions within 100 kb of them (Khyzha *et al*, 2019), which could imply a possible function. The chromosome number, strand, transcript length, exon number and gene length of the DE lincRNAs are presented in Table S3. To locate the mRNAs that were within 100 kb from the lincRNAs, we used Genome Browser online software (https://genome.ucsc.edu/). To find predicted orthologues with high scores of these mRNAs, we utilized the DRSC Integrative Ortholog Prediction Tool (DIOPT)(https://www.flyrnai.org/cgi-bin/DRSC_orthologs.pl) (Table S4). The human mRNA orthologues found included calmegin, NAA15 (also called NMDA receptor-regulated gene 1b) and H2AZ2 (or h2afv, H2A histone family-member V). However, of these genes, only H2AZ2 was DE (padj<0.05, FC>1.5 or <0.67) and then only in the obesity model. Therefore, neighboring mRNAs were not informative as to possible functions of the lincRNAs.

### miRNAs do not seem to link anxiety and obesity in zebrafish larvae

PCA analysis showed distinct miRNA clustering profiles in the obesity and anxiety models (Figures 4A, 4B), differing from those of the poly(A)+ RNAs, although they originated from the same RNA samples. Only one miRNA, dre-miR-738, was DE in the anxiety model (FDR<0.05, fold change>1.5 or < 0.67) (Figure 4C). It was also upregulated in the obesity model, in which 24 other DE miRNAs were observed (FDR<0.05, fold change>1.5 or < 0.67) and this upregulation was validated by qRT-PCR in the obesity but not the anxiety model (Figures 4D, 4F). In short, none of the significant and validated DE miRNA in anxiety were also DE in obesity, contrasting adult mammals, where meta-analysis indicated miRNA participation in the link between anxiety and the metabolic syndrome (Meydan *et al*, 2016).

**Figure 4.**
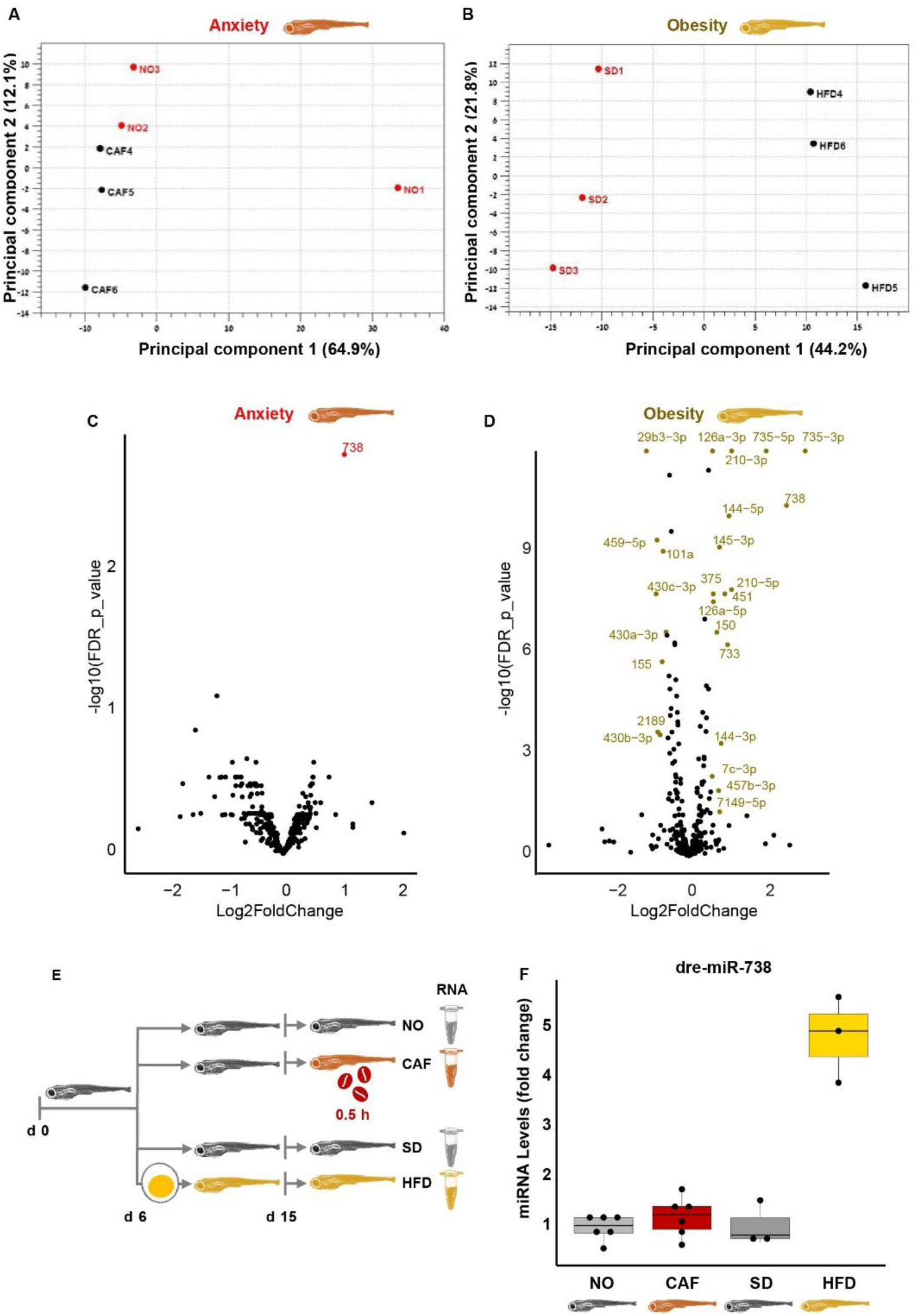
Null involvement of miRNAs in the anxiety and obesity link in zebrafish larvae. A PCA graph of anxiety. B PCA graph of obesity. C Volcano plot of anxiety model. Red points: DE in the anxiety model, FDR <0.05 and absolute log2 fold change>0.58 [>1.5- and <0.67-fold change). The “dre-miR” prefix has been removed from miR labels to enable better visualization. D Volcano plot of obesity model. Gold points: DE in the obesity model, FDR <0.05 and absolute log2 fold change>0.58 [>1.5- and <0.67-fold change). The “dre-miR” prefix has been removed from miR labels to enable better visualization. E Experimental design: zebrafish larvae (6 dpf) were fed either standard diet (SD) or high fat diet (HFD). At age 15 dpf, a group of the standard diet fed larvae were further divided into caffeine-exposed larvae (CAF) and their controls (NO). Following caffeine exposure (0.5 h) of the CAF larvae, larvae of all treatments were snap frozen until RNA extraction for RNA-seq. F qRT-PCR of dre-miR-738 in anxiety and obesity models, Mean ± SE, Student’s t-Test, n = 3-8 (10-30 larvae/replicate, 1-3 independent experiments, where RNA of one of the experiments had been used in RNA-seq. The reference was the average of three miRNAs: dre-miR-let7a, dre-miR-125b-5p and dre-miR-26a-5p.

### Larval tRFs were upregulated in anxiety and downregulated in obesity

Recent studies reveal that tRFs are regulatory tRNA-derived molecules that in certain neuro-pathologies may replace miRNAs in the control of immune system homeostasis (Zorbaz *et al*, 2022; Winek *et al*, 2020). Therefore, DE tRFs were sought. Only 29-32 nts nuclear-derived tRFs were detected. Most DE tRFs were upregulated in anxiety and/or downregulated in obesity, without regard to tRF type (Fig. 5). Thirteen of these tRFs, listed in Table 1, were upregulated in anxiety and downregulated in obesity, similarly to the immune system-related inverse regulation phenomenon observed for poly(A)+ RNAs. Interestingly, three of the inversely regulated tRFs belong to the 5’-half tRF type, which are related to immune system reactions (Zorbaz *et al*, 2022) and have been found to function in neurological disorders (Mathew *et al*, 2022), implying their possible role in anxiety and obesity.

**Fig. 5.**
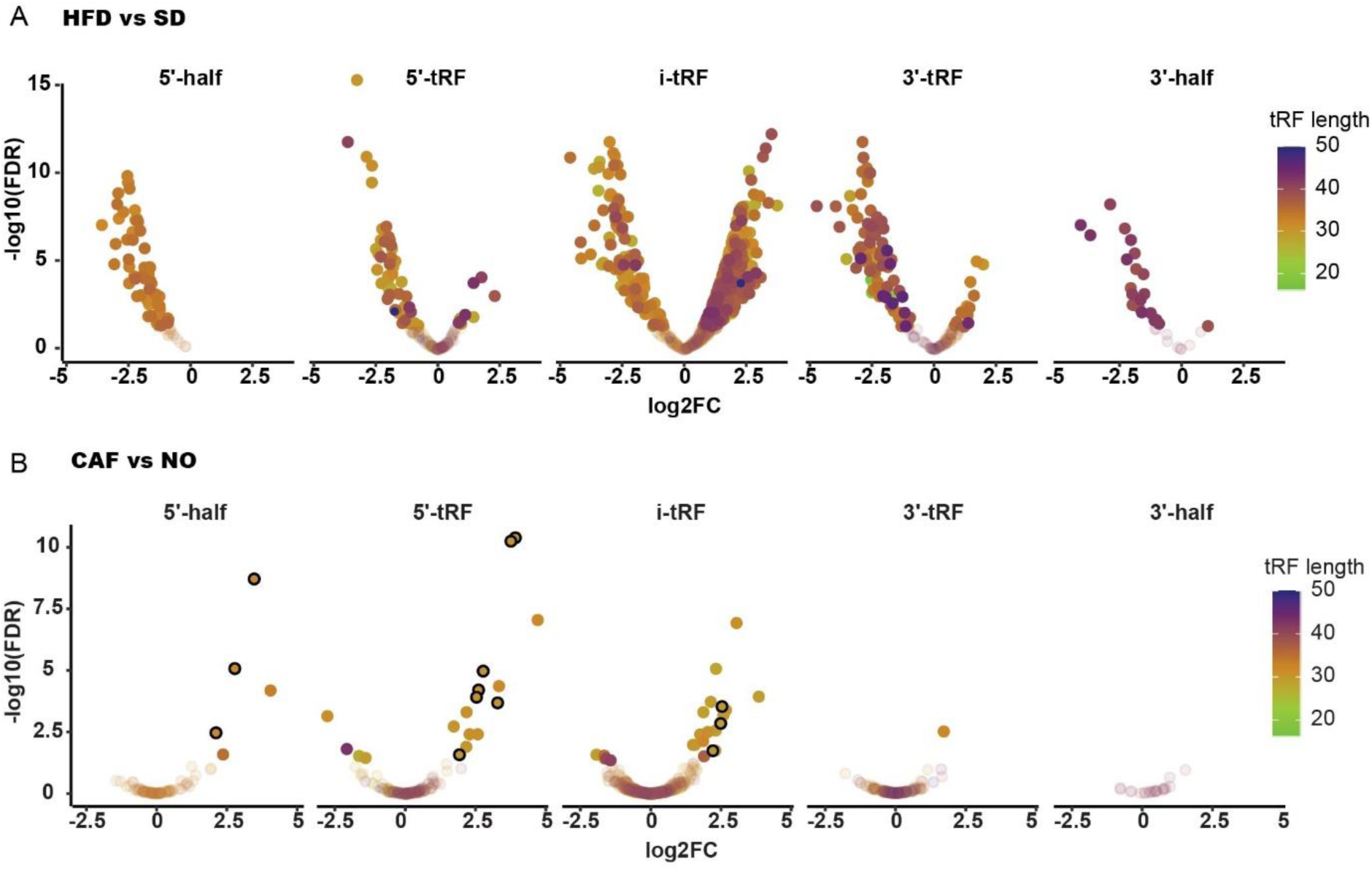
tRFs are also inversely regulated in anxiety and obesity. A HFD vs SD B CAF vs NO - The encircled dots are the 13 tRFs that are upregulated in anxiety and downregulated in obesity.

**Table 1.**
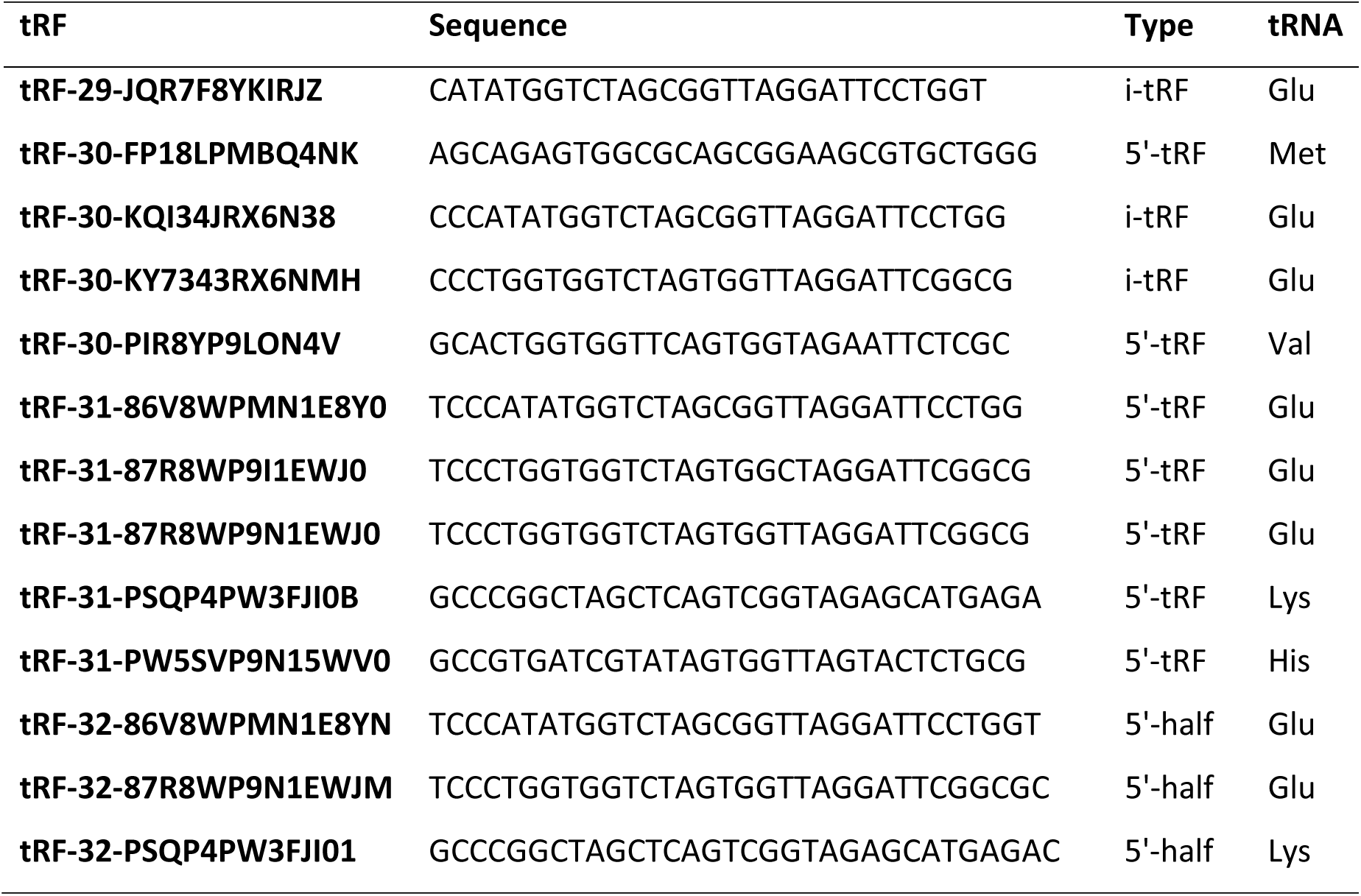
Inversely regulated tRFs - upregulated in anxiety and downregulated in obesity.

### In larvae, obesity fails to accompany anxiety and vice versa

When adult zebrafish become obese, they acquire anxiety-like behavior (Türkoğlu *et al*, 2022; Picolo *et al*, 2021). To examine if this occurs in larvae, we fed larvae a HFD (for 10 days, as in the obesity model) and tested their anxiety-like behavior when they were 15 dpf. No difference in thigmotaxis or erratic swimming was found between SD- and HFD-fed larvae (Figure 6A). Furthermore, neither one (on 7 dpf), two (on 5 and 9 dpf) nor four (on 5, 7, 9 and 12 dpf) high dose caffeine pretreatments (0.5 h, as in the larval anxiety model) increased abdominal adiposity in 15 dpf larvae (Fig. 6B). Unlike adult fish and although larvae have immune-related inverse regulation between anxiety and obesity, larval obesity does not lead to anxiety. The apparent lack of an anxiety-obesity link in zebrafish larvae tentatively indicates that the positive link between anxiety and metabolic disorders in vertebrates may develop with age.

**Figure 6.**
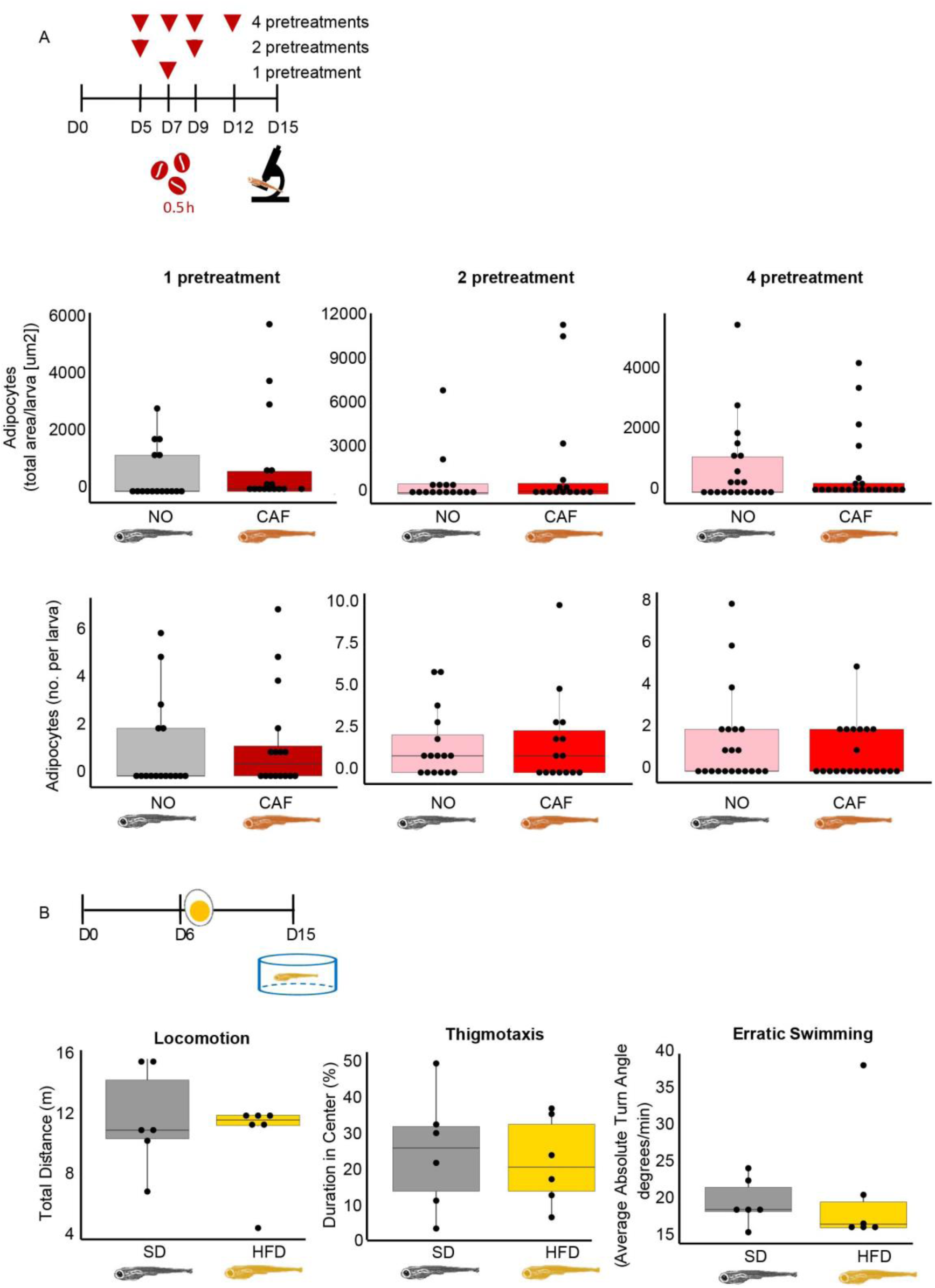
Obese larvae failed to present anxiety-like behavior and vice versa. Presented are the timeline and results demonstrating the effect of: A anxiety (caffeine pretreatments [100 mg/l for 0.5 h]) on obesity (adipocyte size and number in the abdominal area of larvae) B obesity (egg yolk solution-based HFD, 10 days) on anxiety-like behavior (thigmotaxis and erratic swimming). Note the absence of apparent links between the two impacts.

### Age and sex affected mRNA expression

Given the apparently inverse regulation between anxiety and obesity in fish larvae, we tested if this regulation pattern is maintained with age and in both sexes, or if it is particular to the larval developmental stage. For this purpose, we used an accepted anxiety model for adult zebrafish (Wong *et al*, 2010; Rosa *et al*, 2018; Tran *et al*, 2017) to assess the levels of DE transcripts that had been upregulated in our anxiety larval model and downregulated in our obesity larval model. This adult anxiety model was validated in middle-aged adult fish in three independent experiments (each with male and female fish). Notably, caffeine-exposed fish (of both sexes) swam significantly less to the top half of the tank, spent less time in the top half and remained closer to the bottom of the tank, than did those that were not exposed to caffeine (Figure S5). Such behavior is characteristic of anxiety in adult zebrafish (Wong *et al*, 2010; Rosa *et al*, 2018; Tran *et al*, 2017). Results were more pronounced in males. Although both caffeine-exposed females and males crossed to the top half of the tank significantly more than the controls (Figure 7A), only males exposed to caffeine spent significantly more time at the bottom and remained closer to the bottom of the tank than male controls (Figure 7B, C). Despite the clear anxiety-like behavior of the adult zebrafish, of the 6 immune system-related DE transcripts found in larvae, only one, nfkbiab, was DE in adults, increased in the whole-body tissues of females by 24% (p=0.006) and males by 61% (p=0.050) (Figure 7D); this is less than in larvae, in which its level doubled following caffeine exposure. None of the other mRNAs tested were upregulated in whole bodies of adults. Since in a whole-body measurement an increased level of mRNA in one organ may cancel out the decreased level of the same mRNA in another, isolated organs of the above middle-aged fish, i.e., brain, liver, intestine and a segment preceding the tail fin (consisting of skin, muscle and bone), were also assessed for expression of genes that had been upregulated in the anxiety larval model. In contrast to what was observed in the larvae, none of these genes were upregulated in the tested organs of adult fish exposed to caffeine (Figure S6). Rather, nr4a1 was downregulated in the female brain, contrasting with its upregulation in larvae. Ptgs2a (Cox2a) and nr4a1 were not expressed in the livers of these fish. Thus, the upregulation initiated in anxiety in zebrafish larvae did not persist in middle-aged adults, and anxiety-like behavior resulting from caffeine exposure seems to differ among sexes, being more pronounced in adult males than females. This may result from the inherently higher anxiety level in adult zebrafish females compared to males (Fontana *et al*, 2020).

**Figure 7.**
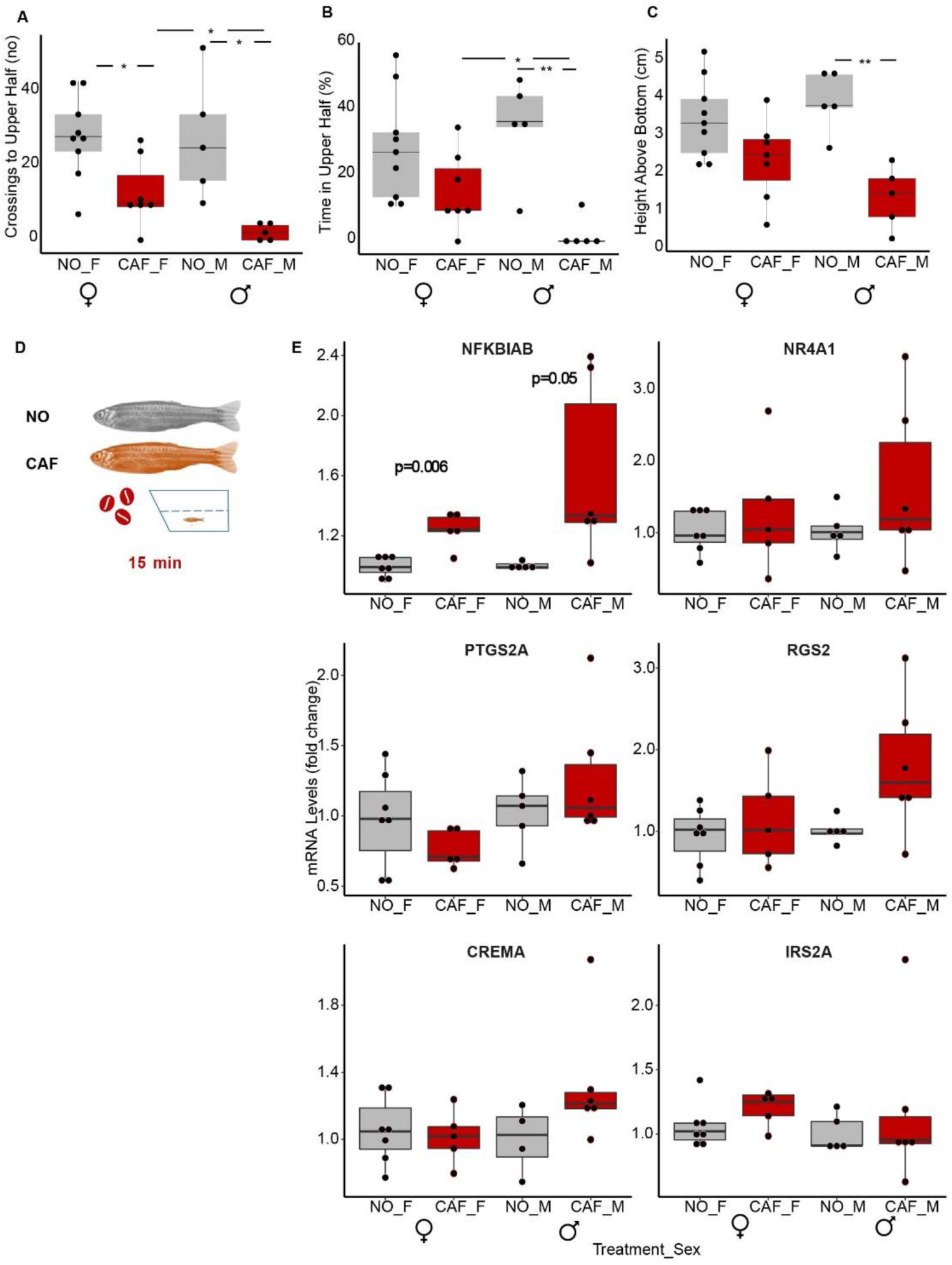
The NFκB inhibitor (nfkbiab) transcript was upregulated in whole bodies of anxiety-induced middle-aged adult males and females. Anxiety-like behavioral tests – A Crossings to upper half of tank (A), B Percent time in upper half of tank (B) C Height above bottom of tank (C) NO F, CAF F – non-treated and caffeine-exposed females, NO M, CAF M – non-treated and caffeine-exposed males. n=5 male and n= 7-9 female fish in 3 independent experiments (each experiment had both males and females). Student’s t-Test, * - p<0.05, ** -p<0.01. D Scheme showing that individual fish were exposed to water or water containing caffeine in a novel diving tank for 15 min, during which movements were recorded. NO – no treatment, CAF – caffeine exposure. E qRT-PCR results - n= 4-6 male fish and 5-7 females in each treatment, 2 independent experiments, with males and females in each. The reference genes were actb2 and eef1a1l1. Student’s t-Test.

### LncRNA profiles change with age and sex

Similar to mRNAs, seven lincRNAs were upregulated by about two-fold and one processed transcript was downregulated by about 30% in the larval anxiety model (mentioned above). However, none of these were up- or downregulated in whole bodies of middle-aged zebrafish with induced anxiety, although we observed higher levels of these lincRNAs in males compared to females. Cautiously, we note that this might be attributed to the eggs in the abdomen of females, which might not express these lincRNAs (Figure S7).

The levels of these seven lincRNAs were also assessed in separate organs of adult male zebrafish, i.e., brain, intestine, liver, testes, and a portion of the tail, consisting of muscle, skin and bone. As found in whole-body middle-aged fish, caffeine had no significant effect on their levels in the various organs (Figure S8). DE was not apparent, conceivably because of a high variation between samples. Moreover, although all seven lincRNAs were found in the muscle/skin/bone sample and in the brain, we could not detect FO681323.202, CABZ01048956-203 and CR926130-201 in the liver and intestine. Disassociation analysis identified two distinct variants of FO834828-201, depending on the tissue; in the muscle, skin and bone samples, it had a melting temperature of ∼75, while in the other organs tested, it had a melting temperature of ∼84. While these lincRNAs also require further analyses, their distinct expression profiles support the notion of age-dependent differences in the links between anxiety and obesity.

### Inversely regulated immune system-related DE transcripts differed between adult zebrafish from larvae

We analyzed two young adult zebrafish RNA-seq datasets, one from whole brains of inherently anxious adult zebrafish (LSB and HSB strains that we termed here SB) versus less anxious ones (AB strain)(Wong & Godwin, 2015; Wong *et al*, 2012), and the other from telencephalons of HFD-versus SD-fed adult zebrafish (that were fed HFD for 11 weeks and tended to be more obese than SD-fed fish) (Meguro *et al*, 2019). The same pipeline used for the larval data was applied to these adult datasets. The individual transcript profiles in both datasets clustered satisfactorily according to treatment (Figures 8A, 8B). There were 3272 anxiety-related DE genes (padj<0.05) compared to only 547 HFD-related DE transcripts (padj<0.05). The larger number of anxiety-DE genes could be strain-related, with only part of the genes associated with anxiety, and the small number of HFD-DE genes attributed to their being brain-derived rather than of liver or adipose tissue sources.

**Figure 8.**
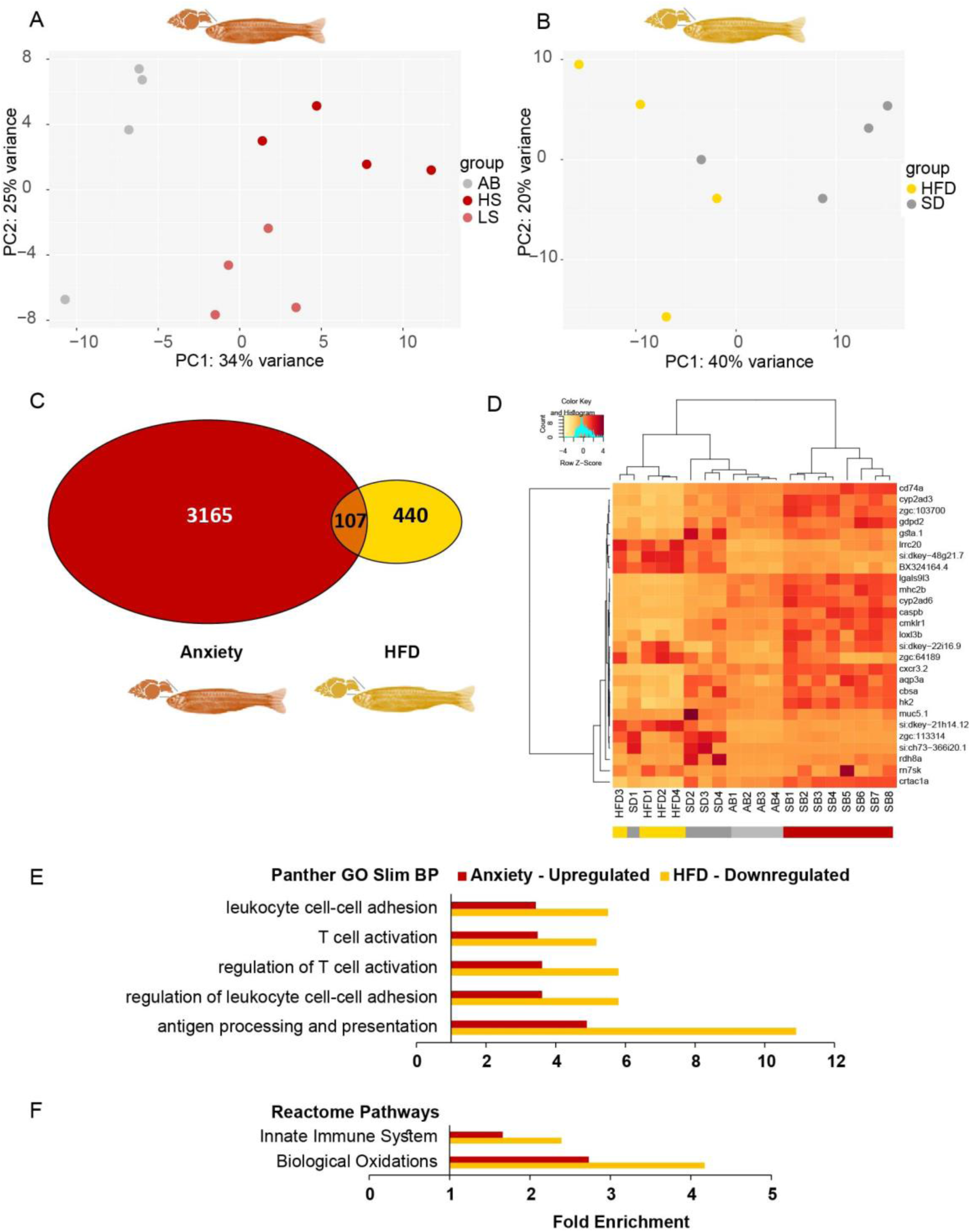
Immune system-related GO terms are enriched among transcripts increased in anxiety and decreased under HFD conditions in adult zebrafish. A PCA of transcripts from young adult zebrafish brains: AB - low anxiety; SB (HS and LS) - high anxiety. B PCA of transcripts from telencephalons of young adult zebrafish fed HFD or SD. C A total of 107 transcripts were commonly DE (padj<0.05) in anxiety (red) and HFD (yellow) in adult zebrafish. D Heatmap of the 27 transcripts upregulated (>1.5) in anxiety among the DE (padj<0.05) intersecting genes in zebrafish adults; AB – Low Anxiety, SB-High Anxiety, SD – Standard Diet, HFD – High Fat Diet. E Overrepresented Panther GO-Slim biological processes (BP) common to upregulated-anxiety and downregulated-HFD genes in zebrafish adults. F Overrepresented Reactome Pathways common to upregulated-anxiety and downregulated-HFD genes in zebrafish adults. Reference transcripts were all found in the Danio rerio database, Fisher’s exact test, correction by FDR<0.05.

As in larvae, more transcripts were upregulated (850, >1.5-fold) than downregulated in anxiety (563, <0.67-fold), and more transcripts were downregulated (378) than upregulated in HFD-fed fish (62). Only 107 transcripts were DE and common to both the anxious and HFD-fed fish (padj<0.05) (Figure 8C), with only two genes appearing also among the 231 anxiety-obesity DE genes in larvae (zgc:162730 and ets2 [v-ets avian erythroblastosis virus E26 oncogene homolog 2]). Most unexpectedly, 20 (74%) of the 27 transcripts upregulated (>1.5-fold) in anxious adult brains, were present among those downregulated (<0.67-fold) in the HFD-fed adult zebrafish (reflected in the heatmap, Figure 8D). Notably, these 20 genes were related to the immune system, e.g., chemokine-like receptor 1 (cmklr1); chemokine [C-X-C motif] receptor 3, tandem duplicate 2 (cxcr3.2); CD74 molecule, major histocompatibility complex, class II invariant chain a (cd74a); caspase b (caspb); lectin, galactoside-binding, soluble, 9 [galectin 9]-like 3 (lgals9l3); and major histocompatibility complex class II integral membrane beta chain gene (mhc2b).

To further assess the function of the DE transcripts in anxious and HFD-fed adult zebrafish, we examined the GO terms of the upregulated and downregulated DE transcripts (padj<0.05) in each condition using Panther. Common GO terms were found only among the upregulated transcripts of the anxious adult zebrafish and the downregulated ones in the HFD-fed adult zebrafish, and these terms were almost exclusively related to the immune system. GO-slim overrepresented biological process (BP) terms included “leukocyte cell-cell adhesion”, “T cell activation” and “antigen processing and presentation”, as well as Reactome Pathways – “innate immune system” and “biological oxidations” (Figures 8E, 8F). Thus, inverse regulation of immune-related genes between anxiety and obesity initiates in larvae and is sustained in adult zebrafish, but is based on different sets of genes, probably since larvae only have the innate immune system, whereas the adults have the innate and adaptive immune systems (Novoa & Figueras, 2012).

## 4. Discussion

Many studies have investigated mechanisms linking anxiety disorders and the metabolic syndrome/obesity (Nousen *et al*, 2014; Xia *et al*, 2021; Mendes *et al*, 2017; Ogrodnik *et al*, 2019; Dutheil *et al*, 2016; Soto *et al*, 2018; Fourrier *et al*, 2019; Pierce *et al*, 2017). However, to our knowledge, none have tested these links in a single study using unbiased transcriptomics to identify the underlying processes. Our findings show that in zebrafish larvae, most of the DE transcripts demonstrated inverse regulation in anxiety and obesity, with anxious larvae presenting mostly upregulated transcripts, contrasting with a downregulation pattern in obese larvae. Moreover, an overrepresented subset of these inversely regulated genes participates in inflammation/immune system-related pathways. While this inverse regulation of immune genes in anxiety-obesity comparisons persisted in adult zebrafish, the genes inversely regulated in the adults differed from those in the larvae. These results raise the questions if this well-known link between anxiety and obesity is subject to age with an antagonistic pleiotropic pattern, and if HFD can protect from anxiety and if so, would such protection persist throughout life.

The long RNA sequencing of the larval anxiety and obesity models revealed that as many as 75% of the genes (including lincRNAs) common to both models that were upregulated in anxiety were also downregulated in obesity, and these inversely regulated DE genes were overrepresented in immune system-related pathways: 1) “CCKR signaling map (gastrin- and cholecystokinin-mediated regulation of cellular processes)”, 2) “inflammation mediated by chemokine and cytokine signaling pathway” and 3) “interleukin signaling pathway”. Correspondingly, downregulation of immune-related genes was revealed in transcriptomes of insulin-resistant zebrafish larvae (Marín-Juez *et al*, 2014).

Apart from the long RNA sequences, thirteen tRFs were discovered to be upregulated in anxiety and downregulated in obesity. These tRFs (29-32 nts) are probably too long to bind to the RISC complex and therefore probably do not act as miRNAs, which are typically ∼22 nts (Pritchard *et al*, 2012); but they may bind certain proteins. Specifically, the inversely regulated tRFs include three 5’ tRF halves that have been implicated in immune related functions (Zorbaz *et al*, 2022) and may potentially participate in immune reactions related to anxiety and obesity. Of these three 5’ halves, tRF-32-PSQP4PW3FJI01, has been found to be upregulated in breast cancer tumors compared to adjacent normal tissue (Wang *et al*, 2019), tRF-32-86V8WPMN1E8YN was upregulated in human malignant mesothelioma vs normal mesothelium (Filetti *et al*, 2022), and tRF-32-87R8WP9N1EWJM has been detected in blood cells (Winek *et al*, 2020). Thus, tRFs may also take part in the immune system-related inverse regulation observed in anxiety and obesity.

One may argue that caffeine exposure or consumption, commonly used to induce anxiety-like behavior in zebrafish (Richendrfer *et al*, 2012; Rosa *et al*, 2018; Tran *et al*, 2017) can be anti-obesogenic. Supporting this notion, in non-obese humans, caffeine consumption (4 or 8 mg/kg body weight, equivalent to 2-3 or 5-6 cups of coffee, respectively) increased the metabolic rate and plasma free fatty acid levels relative to controls (Acheson *et al*, 1980). However, in obese persons on a low calorie diet, caffeine consumption, even relatively high amounts (600 mg/day), has not been effective in promoting weight loss (Dulloo, 2011). In overfed zebrafish larvae, chronic low dose caffeine-exposure reduced the levels of hepatic lipogenic factors and inflammation-related genes and limited hepatic steatosis (Zheng *et al*, 2015). Such low doses improve attention in zebrafish, but do not induce anxiety (Ruiz-Oliveira *et al*, 2019). In contrast, high doses of caffeine generates nervousness in humans (Doepker *et al*, 2016) and anxiety-like behavior in both adult zebrafish (Rosa *et al*, 2018; Tran *et al*, 2017) and zebrafish larvae (Richendrfer *et al*, 2012), compatible with our current findings.

Further study of the link between obesity and anxiety, that was not induced by caffeine, was performed using two adult zebrafish datasets, one from whole brains of inherently anxious adult zebrafish versus less-anxious ones (Wong & Godwin, 2015; Wong *et al*, 2012) and the other from telencephalons of HFD-versus SD-fed adult zebrafish (although the telencephalon of zebrafish differs morphologically from the homologous cerebrum in mammals, it’s mammalian counterparts have similar functions (Friedrich *et al*, 2010)). Surprisingly, as in larvae, 74% of the transcripts upregulated in anxious adults, were present among those downregulated in the HFD-fed adult zebrafish, and these genes included immune-related genes and gene functions, distinct from those found in larvae. In agreement with our findings in anxious adult zebrafish, mice lacking CD4+ T cells did not develop stress-induced anxiety, and CD4+ T cell-secreted xanthine and meningeal T cell derived interferon-γ have been found to participate in murine anxiety-like behavior (Fan *et al*, 2019; Filiano *et al*, 2016). Furthermore, supporting the HFD/obesity-induced downregulation of immune system-related processes found in our study, is the enrichment of the chemokine signaling pathway found among the decreased transcripts in HFD-fed mice cortices (Yoon *et al*, 2019), and the significantly lower weight of HFD-fed rats thymuses compared to controls (McNeilly *et al*, 2015). Moreover, surgery-based weight loss in humans was characterized by enrichment of antigen processing/presentation and interferon γ pathways in subcutaneous fat among the upregulated genes (Poitou *et al*, 2015). Gene ontology terms of one study presented conflicting findings; Stat5a/b knockout-obese mouse hypothalamus upregulated transcripts were associated with insulin stimulus, lipid and saccharide metabolic process, fat cell differentiation and positive regulation of the adaptive and innate immune responses (Fu *et al*, 2017).

Inflammation has been associated with both obesity (Rohm *et al*, 2022; Catrysse & van Loo, 2017; Mraz & Haluzik, 2014) and anxiety disorders (Michopoulos *et al*, 2017; Felger, 2017) and has been studied as a possible link (Nousen *et al*, 2014; Pierce *et al*, 2017; Dutheil *et al*, 2016; Fourrier *et al*, 2019) between both disorders, seeming to contrast our findings of inverse regulation between obesity and anxiety in fish larvae and adults. However, in many of the studies that demonstrated inflammation in both obese and anxious rodents, anti-inflammatory treatments only mitigated anxiety and inflammation, but did not affect obesity. Diet-induced obese mice showed greater anxiety, more inflammation and less insulin signaling in the nucleus accumbens than controls; anti-inflammatory treatment diminished this anxiety and inflammation and improved insulin signaling, but did not reverse HFD-induced obesity (Soto *et al*, 2018). In leptin receptor-null obese mice (db/db vs db/+), chronic food restriction or anti-inflammatory treatment reduced anxiety-like behavior, hippocampal and/or peripheral inflammation, but anti-inflammatory treatment did not reverse the obese state (Fourrier *et al*, 2019). Moreover, acetylcholine (ACh), which is globally elevated in anxiety states (Soreq, 2015), can block NFκB-mediated inflammation (Borovikova *et al*, 2000; Meydan *et al*, 2016); and transgenic excess of miR-132, which exacerbates anxiety (Aten *et al*, 2019) by limiting acetylcholinesterase (which would further reduce inflammation), induces obesity (Hanin *et al*, 2018). Thus, anti-inflammatory treatment can reduce anxiety, but not ameliorate obesity, compatible with our findings of lessened immune-related functions in HFD/obesity. Nevertheless, our results, revealing overrepresentation of immune-related GO terms among the downregulated genes in obese larvae, do not necessarily imply reduced inflammation, since among the genes validated by qRT-PCR (ptgs2a, nfkbiab, crema, rgs2 and nr4a1), most were able to suppress and/or induce inflammation depending on the conditions (Welty *et al*, 2016; Francés *et al*, 2015; Rodríguez-Calvo *et al*, 2017; Catrysse & van Loo, 2017; Cai & Liu, 2012; McNabb *et al*, 2020; Zhu *et al*, 2020). Moreover, the overrepresented immune system related GO terms among the downregulated genes in HFD-fed adult zebrafish belong to the telencephalon of the brain and not to the whole brain, adipose tissue or liver, which if analyzed instead, may have led to a very different conclusion.

Although high fat diet and obesity are often reported to have adverse effects, inducing chronic low-grade inflammation (Mraz & Haluzik, 2014; Poitou *et al*, 2015) and increased risks of diabetes and cardiovascular diseases (Tang *et al*, 2017; Mraz & Haluzik, 2014), recent studies report that HFDs and obesity can have a protective role, in agreement with our study. Indeed, HFD/obesity downregulation of immune function genes could potentially normalize and thus protect zebrafish from exaggerated immune function that can lead to allergies, autoimmune disease and cancer (Provenzano & Deleidi, 2021). For example, 12 weeks of HFD reduced anxiety-like behavior in rats compared to controls (McNeilly *et al*, 2015) and protected mice (fed from 3 weeks-old) from an increased level of anxiety-like behavior that accompanied chronic social stress (Finger *et al*, 2011). Eight weeks of HFD reduced anxiety-like behavior generated by ovariectomy, regardless of the fat type, if it be lard or fish oil (Dornellas *et al*, 2018). In addition, HFD protected C57/Bl male mice from intraperitoneal-injected lipopolysaccharide-induced inflammation in the hypothalamus, although not in the hippocampus (Astiz *et al*, 2016), and also protected against Alzheimer’s disease (AD) pathology in an AD mouse model (Amelianchik *et al*, 2021). HFD-fed offspring of a rat model of type one diabetes, born to dams that were also fed HFD during pregnancy, had a lower incidence of diabetes than offspring fed a normal diet from dams fed a normal diet (Bahr *et al*, 2011). Moreover, twelve months of HFD in transgenic AD mice reduced blood brain barrier leakage and brain atrophy, improved cognition and returned levels of insulin receptor mRNA to that of the normal wild type mice (Elhaik Goldman *et al*, 2018). Furthermore, adult offspring of AD or of wild type mice fed a HFD only during their 3-week gestation, had less memory decline, better synaptic function and less tau (Di Meco *et al*, 2019; Di Meco & Praticò, 2019). Nevertheless, evidence exists that contradicts the positive effect of HFD on cognition in mice and zebrafish (Yoon *et al*, 2019; Meguro *et al*, 2019). Yet, Western diet (high sucrose/HFD)-induced obese minipigs had higher memory scores than lean Western diet controls (Gautier *et al*, 2020). In rats, short term HFD reduced hemorrhagic shock-induced systemic inflammation via CCK receptor stimulation of the afferent vagal nerve and the binding of acetylcholine to nicotinic receptors on macrophages (Luyer *et al*, 2005). In mice, HFD-induced obesity decreased ventilator-induced lung injury, possibly by lowering the levels of matrix metalloproteinase (MMP) activity and soluble receptor for advanced glycation end-products (sRAGE, an epithelial stress marker) in obese mice compared to controls (Wilson *et al*, 2017). In humans, overweightness or obesity were independently associated with lower 60-day in-hospital mortality (Hutagalung *et al*, 2011). Similarly, a meta-analysis and another extensive study (730 intensive care units in 84 countries) showed that obesity was correlated with decreased hospital mortality (Hogue *et al*, 2009). In summary, HFD and obesity can have protective effects. Our current findings in zebrafish hint that these protective effects of HFD and obesity operate via immune pathways, may exert homeostatic effects on the anxiety-upregulated immunity system-associated genes and may relate to the HFD protection reported by others. Understanding this inverse regulation, hence, may aid in further elucidating the mechanisms of metabolic and anxiety disorders.

Based on the above, immune system-related processes are inversely regulated in anxiety and HFD in both zebrafish larvae and adults. However, the specific genes and their functions in larvae differ from those in adults. This raises the speculation that the set of inverse-regulated immune genes in adults may associate with the positive anxiety-obesity link (meaning obesity leads to anxiety and/or vice versa) that has been determined empirically in adult rodents (Dutheil *et al*, 2016; Soto *et al*, 2018; Xia *et al*, 2021), adult humans (Van Reedt Dortland *et al*, 2013; Tang *et al*, 2017; Pierce *et al*, 2017) and adult zebrafish (Türkoğlu *et al*, 2022; Picolo *et al*, 2021). Such a positive anxiety-obesity link was revealed to be lacking in zebrafish larvae, which is in agreement with another study (Türkoğlu *et al*, 2022) and is correlated, as mentioned, with a different set of inversely regulated immune genes than in adult zebrafish. As larvae only have an innate immune system, whereas adult zebrafish have both innate and adaptive immune systems (Novoa & Figueras, 2012), it is conceivable that adaptive immune reactions are important for the development of the positive anxiety-obesity link.

At a more far reaching level, we hypothesize that the development-related link between anxiety and obesity that involves distinct immune system elements at different ages, indicates a possible antagonistic pleiotropic pattern of change in the immune system response. Antagonistic pleiotropy refers to genes that impart traits, which benefit reproduction and fitness under one condition (e.g., young age), but may impede them under another (e.g., older age) (Williams, 1957). The inversely regulated immune system-related genes found in anxiety and obesity in zebrafish may improve survival in young and hinder it later on in life. HFD/obesity can be advantageous when food is scarce and for young larvae, which may have a high daily energy expenditure, if similar to human infants (Pontzer *et al*, 2021). However, extreme anxiety or maternal anxiety disorders is harmful (Misri & Kendrick, 2007). The inversely regulated anxiety-obesity immune system genes are compatible with the lack of an anxiety-obesity link during early development, i.e., obese larvae will not develop anxiety, conferring a survival advantage, whereas in adult zebrafish, the anxiety-obesity immune system correlates with an inversely regulated anxiety-obesity link, i.e., obesity leads to anxiety (Türkoğlu *et al*, 2022; Picolo *et al*, 2021), which is disadvantageous. This link may be due to the need to avoid disrupted physiological homeostasis, causing or preventing age-related diseases. Therefore, the immune system-based link between anxiety and obesity may have an antagonistic pleiotropic pattern.

## Materials and Methods

### Experimental Design

Two zebrafish larval models, one of anxiety (induced by caffeine exposure) and the other of obesity (induced by HFD) were optimized and validated, the anxiety model by behavioral and cortisol level tests and the obesity model by histology-based adiposity analysis. Subsequently, RNA was extracted, sequenced and analyzed to identify and examine the genes that were commonly DE in both models. In addition, the effect of obesity on anxiety, and vice versa, was investigated in zebrafish larvae. The larval RNA-sequencing and qRT-PCR results were compared with data analyzed from adult zebrafish, comprising qRT-PCR results of caffeine-induced anxiety in middle-aged adult zebrafish and RNA-sequencing analyses of two datasets: inherently anxious versus less anxious strains and HFD-fed versus standard diet (SD)-fed young adult zebrafish, using the same pipeline as employed for the zebrafish larval models. Experiments with zebrafish larvae and adults were carried out under the approval of the National Council for Experiments in Animals, numbers IL-19-11-447 and IL-20-10-458, respectively.

### Larvae and Adult Fish

Wild-type zebrafish (AB strain) parents were kept under a 14 h light/10 h dark regimen with lights turned on at 8:00 am. Fish water conditions were pH 7-7.7, 27.7-28.3 °C and 490-510 µS/cm conductivity. Parent fish were fed live 24-hour-old artemia five mornings a week, and standard dry zebrafish food (Gemma micro 300, Skretting, USA) five evenings a week and one morning a week. Fish were not fed on the seventh day. Eggs were gathered and placed in Petri dishes (100 mm diameter) at a density of less than 100 per 30 ml fish water, containing methylene blue (0.00003%). When larvae were 6 days post fertilization (dpf), they were transferred to 2-liter tanks containing 1 liter of fish water (up to 100 larvae per liter of water, up to 4.5 cm deep) and fed a SD (Gemma micro 75, Skretting, USA), twice a day, five times a week, and once a day on the other two days, until 15 dpf (in behavior testing, RNA extraction or cortisol testing) or 16 dpf (in adiposity assessment).

For the larval obesity model, larvae were fed a HFD, based on 20 ml hard-boiled egg yolk solution in addition to the SD. Controls were only fed the SD. The hard-boiled egg yolk solution (den Broeder *et al*, 2017) is based on yolks from chicken eggs bought fresh and boiled for 5 min. The egg yolks were mashed and added to fish water (ratio of 1 g/15 ml). After shaking the mixture vigorously for 5 min, large particles were allowed to settle out during one hour, and the egg yolk solution was divided into aliquots and frozen at -20 °C. The tanks containing the larvae were cleaned regularly and kept under the same light and temperature conditions as those of the parent fish. Zebrafish larvae were euthanized by immersion in ice cold water for at least 30 min (Van Den Bos *et al*, 2017).

Adult fish (15-20 months post fertilization [mpf]) with decreased reproductive activity (i.e., produced less eggs) were used for the adult fish experiments. They were kept under the same conditions as the parent fish (described above) or at a density of less than 16 zebrafish in 3 liters of water in tanks that were cleaned regularly. Euthanization was by gentle transfer and immersion in ice cold water for at least 30 min (Zang *et al*, 2019; Serikuly *et al*, 2020).

### Caffeine Exposure

Prior to caffeine exposure, we exposed larvae to a one-hour habituation period at 28 °C under light. Subsequently, larvae were exposed to caffeine (Sigma, Israel), which induces anxiety-like behavior both in zebrafish larvae (Richendrfer *et al*, 2012) and adult fish (Rosa *et al*, 2018; Tran *et al*, 2017). Caffeine was added as a concentrated solution to the water (final concentration 100 mg/l) containing larva(e) in wells of plates for 0.5 h at 28 °C between 9:00 am - 12:30 pm. In behavior tests, we continued caffeine exposure during the tests. In all other assays, groups of larvae, which were held in a Falcon® 70 µm cell strainer (Corning, USA) in a well of a plate, were transferred with the strainer immediately following the caffeine exposure period to ice cold water for immobilization. Afterwards, excess water was removed, and larvae were flash frozen in liquid nitrogen. Control larvae of this model were manipulated in the same manner as the caffeine-exposed larvae, but without caffeine exposure. Adult fish were exposed to caffeine (100 mg/l) for 15 min (before 1:00 pm), the acceptable duration for inducing anxiety-like behavior in adult zebrafish (Rosa *et al*, 2018; Tran *et al*, 2017). Controls were treated the same as caffeine-exposed adult fish, but without caffeine exposure.

### Larval Behavior Tests

Behavior tests were carried out to verify that caffeine induced anxiety-like behavior in larvae, as measured by thigmotaxis (the preference for the border) and erratic swimming (increased average of cumulative absolute turn angles/min). Thigmotaxis is a particularly exact measure of anxiety, as it is induced by anxiogenic drugs and ameliorated by anxiolytic treatments (Richendrfer *et al*, 2012). Larvae were distributed, one larva per well (wells of 12-well plates contained 2 ml fish water). Following habituation and caffeine exposure, the plates were transferred to the illuminated (intensity = 100%) compartment of the DanioVision instrument in the laboratory. After an additional 10 min habituation period, larval movements were videoed and recorded for 1 h. EthoVision XT 11.5 software (Noldus Information Technology, Wageningen, The Netherlands) was used for analysis, and the parameters measured were distance moved (mm), time duration in the center relative to the whole area (center diameter = 16.3 mm, whole area diameter = 24.1, leaving a peripheral width of 3.8 mm) and turn angle (culmative absolute degree changes/min). The center diameter was chosen, since larvae at 15 dpf are approximately 0.5 mm long (Singleman & Holtzman, 2014), and the location of the larva was determined according to their center points.

### Adult Fish Behavior Tests

The novel diving tank (10.5 cm depth X 27 cm top length X 22 cm bottom length X 16 cm height) contained 2 liters of water at a depth of 10 cm (top half of the tank = upper 5 cm layer of water). Water temperatures during the test were either 25 °C - 28 °C or 21 °C - 23 °C. Although the accepted temperature for adult zebrafish is 24 °C – 29 °C (Aleström *et al*, 2019), their behavior at the lower temperature did not differ from that at the higher temperature. All behavior tests were carried out between 9:00 am and 1:00 pm. The novel diving tank was positioned inside a box (height – 47 cm, width – 51 cm and depth – 38 cm) that was open from one side, reducing distractions for the fish. The tank was illuminated from behind (by an A4 Graphic Tablets LED Light Box Pad Electronic USB Tracing Art Copy Board, Game Fanatics’ Store) for better resolution. Each fish was gently transferred to the novel diving tank containing water, with or without caffeine and videoed individually for 15 min (by a Digital Color Camera) and Debut Video Capture and Screen Recorder Software (NCH Software), which recorded the longitudinal swimming behavior (25 frames per second). The resulting videos were translated into data by a custom MATLAB script. The last 6 min of the fish behavior tests were used to evaluate anxiety-like behavior according to the parameters of height above tank bottom, percent time in the upper half of the tank and the number of times fish crossed to the upper half of the tank.

### Larval Whole-Body Cortisol Assay

Whole-body cortisol levels were determined in larvae that had or had not been exposed to caffeine (pools of 30 larvae per replicate, 15 larvae per well of 6-well plates (containing Falcon® 70 µm cell strainer [Corning, USA] and 10 ml fish water). Following caffeine exposure, larvae were transferred with the strainer to ice cold water, which also prevents any further increase in cortisol (Yeh *et al*, 2013). Cortisol was extracted according to a published procedure (Yeh *et al*, 2013). Briefly, 150 µl water was added to the larval pools, for homogenization by a motorized pestle and a minimum of ten subsequent passages through a syringe needle (20-G). Ethyl acetate (1 ml) was added, followed by vortexing (30 sec) and centrifugation (5 min, 3000 X G, 4 °C). After freezing the aqueous layer, the solvent was transferred to a new tube and evaporated at 30 °C under nitrogen. The cortisol that remained was dissolved in phosphate-buffered saline -0.2% bovine serum albumin (to be divided between two technical replicates) and frozen. Before continuing with an enzyme-linked immunosorbent assay (ELISA) test in a Cortisol Parameter Assay Kit (R&D Systems, USA), samples were vortexed on a thermomixer for 5 min at 1100 rpm at 37 °C.

### Adipocyte Assessment Assay

Larvae were washed and subsequently stained with Nile Red (Sigma, Israel), which specifically stains neutral lipids, such as triglycerides (Minchin & Rawls, 2011). Larvae were exposed to Nile Red (500 ng/ml) for 0.5 h, and subsequently euthanized. Using 1% low melting point agarose or a glycerol-based mounting medium (ibidi Mounting Medium, ibidi, Germany), each larva was positioned on its right side (location of adipocyte deposit) on a coverslip assembly, based on a published diagram (Westerfield, 2007) (wiki.zfin.org/display/prot/Viewing+Chambers). The degree of obesity was assessed by the number of Nile Red-stained lipid droplets (assumed to be in adipocytes) and their area, as determined by confocal microscopy (Zeiss LSM 700 laser scanning confocal microscope, using a X5 objective, laser settings of 488 nm for excitation and ≥539 nm for emission and ZEN lite 2012 software (Zeiss)). Conditions, including the laser power, pinhole, master gain and digital offset were kept constant for each experiment. The layer counted was that with the most Nile Red-stained lipid droplets in the abdominal area. Lipid droplets smaller than 100 µm2 were excluded, as were those appearing in a row (since they could be fat droplets in lymph vessels and not adipocytes).

### Caffeine Pretreatments

In experiments examining the effect of anxiety on obesity, larvae were pretreated with caffeine, and assayed for the accumulation of adipocytes on day 15 post fertilization (pf), as described above. The time of day, concentration and duration of exposure to caffeine were the same as for the other assays, but treatments were done either on day 7, days 5 and 9 or days 5, 7, 9 and 12 pf, in the one, two and four caffeine-pretreatment experiments, respectively. For pretreatments, larvae (up to 25) were gently concentrated to 50 ml water, to which was added or not added a concentrated solution of caffeine. Following the half hour exposure to caffeine (0 or 100 mg/l), larvae were gently transferred to their original tank using a 150 µm filter.

### RNA Extraction

RNA was isolated from 1) pools of larvae and 2) whole bodies or individual organs of adult fish. All extractions were preceded by homogenization using a motorized pestle and at least 10 subsequent passages through a syringe needle (20-G). Extraction from adult whole bodies (minced) or a small portion of the tail area (skin, muscle and bone) was by QIAzol Lysis Reagent according to the Qiazol Protocol (Qiagen, Germany); from livers and intestines of the adult fish, using Tissue Total RNA Mini Kit (Geneaid Biotech Ltd, Taiwan); and from larvae and brains, by the Nucleospin® miRNA kit (Macherey-Nagel GmbH & Co. KG, Germany). All extractions were according to the manufacturer’s recommendations.

### RNA-seq

Twelve RNA samples were sequenced. Each larval sample was a biological replicate derived from a pool of 30 larvae (15 dpf). There were three replicates per treatment and four treatments: no treatment (NO) and caffeine (CAF) of the anxiety model and standard diet (SD) and high fat diet (HFD) of the obesity model. The CAF samples were larvae that had been exposed to caffeine in groups of 15 larvae per well of 6-well plates (containing Falcon® 70 µm cell strainer (Corning, USA) and 10 ml fish water) and two wells per sample. The NO samples were the controls of the CAF samples and therefore were manipulated in the same manner, but with no treatment. The SD samples were the controls of the HFD fed larvae. The larvae of these samples were fed either SD or HFD from 6 dpf until 15 dpf. Subsequently, they were collected, immobilized and flash frozen in the same manner as the NO and CAF larvae. Samples were kept at -80 °C until RNA extraction.

Integrity assessment with an Agilent TapeStation, library preparation and sequencing of the RNA were carried out at the Technion Genomics Center, Technion-Israel Institute of Technology. Samples had an RNA integrity number (RIN) > 8.5, except for the HFD samples, with RINs of 7.8-8.2, possibly because of the high fat content. Twelve poly(A)+ and 12 short RNA libraries were prepared. The short RNA libraries were generated using QIAseq miRNA Library Kit, and the poly(A)+ RNA libraries were generated using TruSeq RNA Library Prep Kit v2, both according to manufacturer’s protocol. Sequencing was on Illumina HiSeq 2500, generating 75 nucleotides single-end runs from the short RNA libraries and 51 nucleotides single-end runs from the poly(A)+ RNA libraries.

Poly(A)+ analysis - The poly(A)+ RNA libraries yielded from 14,745,028 to 20,885,465 reads per sample. As assessed using FastQC, version v0.11.9, the per base sequence quality was >32, except for base 5, in all sequences. None of the sequences had overrepresented sequences or adapter content. Trimming (Trimmomatic v. 0.36) was not carried out, since it did not improve sequence quality, according to FastQC. The zebrafish reference genome used for mapping was Danio_rerio.GRCz11.dna.primary_assembly.fa.gz (the primary assembly of the genome, which contains all top level genes. Haplotypes [README from ftp://ftp.ensembl.org/pub/release-98/fasta/danio_rerio/dna/]) were excluded, to simplify analysis. For annotation, Danio_rerio.GRCz11.98.gtf.gz was used, which includes both genes with chromosomal locations and genes whose chromosomal locations have yet to be determined (README from ftp://ftp.ensembl.org/pub/release-98/gtf/danio_rerio/). We used RSEM software for read alignment (BOWTIE2), annotation and expression calculation (https://bioinformaticshome.com/tools/rna-seq/descriptions/RSEM.html). The RSEM model statistics, obtained by RSEM for the 12 samples, showed that the fragment lengths were similar (51 ± 0.7, mean ± sd), the read lengths were identical (51 ± 0), observed quality vs Phred Quality score graphs were similar, and the alignment statistics were also similar (48-50% multialigned, 34-36% unique, 15-18% unalignable).

DE analysis of RNA isoforms was carried out by the RSEM wrapper of EBSeq. An ngvector was generated to group information, which is necessary for EBSeq isoform-level differential expression. The resulting EBSeq output was a matrix with normalized mean counts for each transcript of each treatment. Those genes that had a false discovery rate (FDR) < 0.05 were obtained by the RSEM wrapper of FDR. In EBSeq, there are 15 patterns from which one can determine DE patterns, relevant for determining isoforms that were DE in both the anxiety model and the obesity model. Pattern 4 – Caf and HFD are the same but different from SD and NO, which are the same; Pattern 9 – Caf and HFD are the same but different from SD and NO, which are different; Pattern 14 – Caf and HFD differ, but SD and NO are the same; and Pattern 15 – all treatments differ. These pattern subsets were determined and fold changes were calculated. Isoforms were considered to be DE if the fold change had an absolute log2 > 0.58 (in real numbers - >1.5 or <0.67) and a FDR<0.05.

In addition, mRNA transcriptomic analysis was carried out using DESeq2 (https://bioconductor.org/packages/release/bioc/html/DESeq2.html) of Bioconductor 3.10 (https://www.bioconductor.org/install/) and R software 3.6.1 (https://www.r-project.org/). Genes were considered to be DE if they had a significance of p<0.05 and a fold change >1.5 fold or < 0.67. Gene functions were analyzed by the overrepresentation test of Panther Gene Ontology, version 17.0 (http://pantherdb.org/geneListAnalysis.do); those used were Panther GO-slim biological process (BP), Panther GO-slim molecular functions (MF), Panther GO-slim cellular components (CC), Panther Pathways, Reactome Pathways and Panther Protein Class. In these tests, genes are considered to be overrepresented if they appear significantly more than in the Danio rerio reference genome (and they are considered to be underrepresented, if they appear significantly less). The following gene groups were tested as pairs to find the GO-terms and pathways that were commonly overrepresented in both: Upregulated (FC>0, padj<0.05) anxiety and obesity DE gene lists; upregulated anxiety DE genes with downregulated (FC<0, padj<0.05) obesity DE genes; and downregulated anxiety DE genes with each of these two obesity DE gene lists. The RNA-seq poly(A)+ results have been deposited at the database repository Gene Expression Omnibus (GEO), as SuperSeries GSE207550/SubSeries GSE207457.

Short RNA analysis (miRNAs and tRFs) - Short RNA quality was analyzed by the Technion Genomics Center. FastQC version 0.11.5 demonstrated that the reads had a high quality, i.e., Phred Score > 33 and 26,126,096 to 38,085,377 reads per sample.

For miRNA analysis, the reads were trimmed using the online CLC Genomics Workbench 20.0.3, which included removal of the Qiagen 3’ adapter on the 3’ end. Reads without this adapter were discarded, and only sequences between 15 and 54 nucleotides were retained. These trimmed files (read lengths of ∼23 bases) were joined. miR quantify was run on these files, using the reference - miRbase v.22.1 and the preference - Danio rerio. Normalization was executed using TMM (trimmed mean of M values; TMM normalization adjusts library sizes based on the assumption that most genes are not DE). Annotation was carried out using Go Mapping RNAcentral v10. The miRNA-seq results have been deposited at GEO, as SuperSeries GSE207550/SubSeries GSE207549.

For tRF analysis, the short RNA sequencing was aligned to tRFs using MINT-map (Loher *et al*, 2017), according to the default settings of the pipeline. All tRFs for which the 75th quantile was lower than the mean over all samples were removed from the count data a priori (13 tRFs excluded). Differential expression of tRFs was assessed using the edgeR package (Robinson *et al*, 2009) and graphs were plotted using ggplot2 (Wickham, 2016).

### cDNA Synthesis

For mRNA and lncRNA - The Verso cDNA Synthesis Kit (ThermoFisher Scientific, USA) was employed to synthesize cDNA for qRT-PCR from the RNA extracted above. It was carried out based on the manufacturer’s recommendations. Briefly, RNA (0.25 -1 µg) was heated to 70 °C for 5 min and cooled to 4 °C, before adding the anchored-oligo dT primer and other reagents supplied in the kit, including the RT enhancer that prevents genomic DNA carryover. The cDNA synthesis reaction was executed at 42 °C for 60 min, followed by 52 °C for 30 min, and lastly, enzyme inactivation at 95 °C for 2 min.

For miRNAs, cDNA was synthesized based on a published protocol (Balcells *et al*, 2011). Briefly, the 10 µl reaction, which included 1 µg RNA, 1 µl of 10X poly(A) polymerase buffer (New England Biolabs, USA), 0.25 mM ATP, 1 mM dNTPs (New England Biolabs, USA), 1 µM RT-primer (GGTCCAGTTTT TTTTTTTTTTTVN [V is A, C and G; N is A, C, G and T]), 80 U MuLV reverse transcriptase (New England Biolabs, USA) and 0.75 U poly(A) polymerase (New England Biolabs, USA), was subjected to 42 °C for 1 h, followed by enzyme inactivation at 95 °C for 5 min.

### Quantitative Reverse Transcription - Polymerase Chain Reaction [qRT-PCR])

cDNA was amplified in a reaction that contained the iTAQ Universal SYBR Green Supermix (Bio-Rad, Israel) (based on the protocol supplied by the manufacturers) and specific primers (Table S1) in the 7900HT Applied Biosystems instrument. Primers (Merck, Israel) for mRNA and lncRNA were designed using the NCBI Primer-Blast software, and primers for miRs were designed according to Busk (Busk, 2014). For normalization in mRNA and lncRNA qRT-PCR, the average of the reference genes, Danio rerio eukaryotic translation elongation factor 1 alpha 1, like 1 (eef1a1l1, NM_131263.1) and Danio rerio actin, beta 2 (actb2, NM_181601.5), was used. These genes have been reported to be good references for qRT-PCR in zebrafish (Tang *et al*, 2007). For miR qRT-PCR, the average of the three miRs, dre-miR-let7a, dre-miR-125b-5p and dre-miR-26a-5p, was employed for normalization. These miRs were chosen, since their absolute fold change was ≤ 1.1 according small RNA-seq analysis using CLC Genomics Workbench v.20. The PCR cycles were as follows: denaturation at 95 °C for 1 min followed by 40 cycles of 95 °C for 15 s and 58 °C for 30 s. Analysis was according to the 2 ^-ΔΔCt^ method.

### Datasets

#### Brains of anxious vs less anxious adult zebrafish -

Transcriptomic analysis of brain samples from a publicly available dataset (accession number GSE61108) of adult zebrafish with different inherent levels of anxiety-like behavior was carried out. The strains in the dataset included AB (also used in the above experiments), which is a low anxiety-like strain, common in laboratories, and two other strains with higher anxiety-like behavior, termed here collectively as SB, comprising LSB (low stationary behavior) and HSB (high stationary behavior) (Wong & Godwin, 2015; Wong *et al*, 2012). Each strain was represented by four pooled samples of 10 whole brains (two males and two females); thus, for our analysis there were 4 samples of AB and 8 samples of SB (4 HSB and 4 LSB). cDNA libraries had been generated with TruSeq RNA Sample Prep V2 of Illumina and sequenced as 72 bp single ends on Illumina (GA_IIX_), similarly to our procedures. To analyze this dataset, we used a similar procedure to the RNA analysis for larval samples, with one exception. FastQC, version v0.11.9 revealed TruSeq adapters, which we removed with Trimmomatic v.0.36 and further confirmed removal by FastQC analysis. The remaining analysis was executed with RSEM and DESeq2, as described above.

#### Telencephalons of HFD-fed vs SD-fed adult zebrafish -

A dataset of transcripts from 3 months post fertilization male zebrafish telencephalons, as described in the article (Meguro *et al*, 2019), was kindly provided by Dr. Shinichi Meguro from the Kao Corporation, Japan. In brief, the fish were fed an HFD of 80% control diet and 20% lard (w/w) or the SD (control diet) for 11 weeks. Euthanization was by immersion in tricaine for the dissection of the telencephalon. The libraries had been prepared using the TruSeq RNA Sample Preparation Kit v2 of Illumina and sequenced on the HiSeq 2500 platform (Illumina Inc) as 101 bp paired ends. We used FastQC, version v0.11.9 to analyze read quality and Trimmomatic v.0.36 to remove any adapters. Their removal was confirmed by FastQC analysis. The remaining analysis was executed with RSEM and DESeq2, as described above.

### Statistical Analysis

Student’s t test was employed for statistical analysis, using the two-tailed test, unless noted otherwise. Results were considered to be statistically significant if p<0.05.

## Acknowledgments

Special thanks to Dr. Shinichi Meguro from the Kao Corporation, Japan for sharing his dataset with us, to Dr. Karen Jackson for the use of zebrafish equipment and instruments and to Dr. Tamara Zorbaz and Ms. Petra Pollins for indispensable help with the figures. This study was supported by the Israel Science Foundation award [grant number 1016/18] to H.S. and an internal MIGAL grant to A.M.

## Author Contributions

**Hila Yehuda**: Conceptualization; data curation; validation; formal analysis; investigation; methodology; writing – original draft: writing – review and editing. **Nimrod Madrer:** investigation; formal analysis; writing – review and editing. **Doron Goldberg:** software; writing – review and editing. **Ari Meerson**: Conceptualization; supervision; funding acquisition; writing – review and editing. **Hermona Soreq:** Conceptualization; supervision; funding acquisition; writing – original draft: writing-review and editing.

## Conflict of interest

The authors declare that they have no conflict of interest.

## The Paper Explained

### Problem

Anxiety disorders have been linked to metabolic disorders, such as obesity, in humans, rodents and adult fish, but the mechanisms behind the link are still unclear. Further elucidation of such a link could aid in prevention or therapy of these two disorders. One way to examine the link between anxiety and obesity is to identify genes that are upregulated or downregulated in both disorders by executing a single RNA-sequencing study. To our knowledge this has yet to be done.

### Results

According to gene expression comparisons in anxiety and obesity zebrafish larval models, as well as in inherently anxious versus less anxious adult zebrafish strains and high fat diet-versus standard diet-fed adult zebrafish, most genes (including regulatory noncoding RNAs e.g., long noncoding RNAs and transfer RNA fragments) upregulated in anxiety were downregulated in obesity in both zebrafish larvae and adults. Many of the genes were related to immune system processes and pathways that differed between larvae (in which obesity did not lead to anxiety or vice versa) and adult zebrafish (in which obesity is known to lead to anxiety).

### Impact

HFD downregulation of genes that are upregulated in anxiety may explain HFD protective effects that have been reported for anxiety and other neurological disorders in animal models. Moreover, the age-related adjustment in immune system regulation and its relation to the anxiety-obesity link may improve the understanding of these two disorders.

## For More Information

https://bioinformaticshome.com/tools/rna-seq/descriptions/RSEM.html

https://bioconductor.org/packages/release/bioc/html/DESeq2.html

https://www.bioconductor.org/install/

https://www.r-project.org/

http://pantherdb.org/geneListAnalysis.do/

https://genome.ucsc.edu

https://www.flyrnai.org/cgi-bin/DRSC_orthologs.pl/

## Data Availability Section

The RNA-seq data used in this study have been depositied in the Gene Expression Omnibus (GEO). The RNA-seq poly(A)+ dataset has been deposited as SuperSeries GSE207550/SubSeries GSE207457. The small RNA-seq dataset has been deposited as SuperSeries GSE207550/SubSeries GSE207549.

## Notes

### Competing Interest Statement

The authors have declared no competing interest.

## References

de Abreu MS, Giacomini ACVV, Zanandrea R, dos Santos BE, Genario R, de Oliveira GG, Friend AJ, Amstislavskaya TG & Kalueff A V. (2018) Psychoneuroimmunology and immunopsychiatry of zebrafish. Psychoneuroendocrinology 92: 1–12

Acheson KJ, Zahorska-Markiewicz B, Pittet P, Anantharaman K & Jéquier E (1980) Caffeine and coffee: Their influence on metabolic rate and substrate utilization in normal weight and obese individuals. Am J Clin Nutr 33: 989–997

Aleström P, D’Angelo L, J. P, Schorderet DF, Schulte-Merker S, Sohm F & Warner S (2019) Zebrafish: Housing and husbandry recommendations. Lab Anim 54: 213–224

Almeida-Suhett CP, Graham A, Chen Y & Deuster P (2017) Behavioral changes in male mice fed a high-fat diet are associated with IL-1β expression in specific brain regions. Physiol Behav 169: 130–140

Alonso-Caraballo Y, Hodgson KJ, Morgan SA, Ferrario CR & Vollbrecht PJ (2019) Enhanced anxiety-like behavior emerges with weight gain in male and female obesity-susceptible rats. Behav Brain Res 360: 81–93

Amelianchik A, Merkel J, Palanisamy P, Kaneki S, Hyatt E & Norris EH (2021) The protective effect of early dietary fat consumption on Alzheimer’s disease–related pathology and cognitive function in mice. Alzheimer’s Dement Transl Res Clin Interv 7: 1–11

Aprile M, Katopodi V, Leucci E & Costa V (2020) Lncrnas in cancer: From garbage to junk. Cancers (Basel) 12: 1–32

Arner P & Kulyté A (2015) MicroRNA regulatory networks in human adipose tissue and obesity. Nat Rev Endocrinol 11: 276–288

Aryal B, Singh AK, Rotllan N, Price N & Fernández-Hernando C (2017) MicroRNAs and lipid metabolism. Curr Opin Lipidol 28: 273–280

Astiz M, Pernía O, Barrios V, Garcia-Segura LM & Diz-Chaves Y (2016) Short-term high-fat diet feeding provides hypothalamic but not hippocampal protection against acute infection in male mice. Neuroendocrinology 104: 40–50

Aten S, Page CE, Kalidindi A, Wheaton K, Niraula A, Godbout JP, Hoyt KR & Obrietan K (2019) miR-132/212 is induced by stress and its dysregulation triggers anxiety-related behavior. Neuropharmacology 144: 256–270

Bahr J, Klöting N, Wilke B, Klöting I & Follak N (2011) High-fat diet protects BB/OK rats from developing type 1 diabetes. Diabetes Metab Res Rev 27: 552–556

Baker KD, Loughman A, Spencer SJ & Reichelt AC (2017) The impact of obesity and hypercaloric diet consumption on anxiety and emotional behavior across the lifespan. Neurosci Biobehav Rev 83: 173–182

Balcells I, Cirera S & Busk PK (2011) Specific and sensitive quantitative RT-PCR of miRNAs with DNA primers. BMC Biotechnol 11: 70

Blay SL & Marinho V (2012) Anxiety disorders in old age. Curr Opin Psychiatry 25: 462–467

Borovikova LV., Ivanova S, Zhang M, Yang H, Botchkina GI, Watkins LR, Wang H, Abumrad N, Eaton JW & Tracey KJ (2000) Vagus nerve stimulation attenuates the systemic inflammatory response to endotoxin. Nature 405: 458–462

Van Den Bos R, Mes W, Galligani P, Heil A, Zethof J, Flik G & Gorissen M (2017) Further characterisation of differences between TL and AB Zebrafish (Danio rerio): Gene expression, physiology and behaviour at day 5 of the larval stage. PLoS One 12: 1–15

den Broeder M, Moester M, Kamstra J, Cenijn P, Davidoiu V, Kamminga L, Ariese F, de Boer J & Legler J (2017) Altered Adipogenesis in Zebrafish Larvae Following High Fat Diet and Chemical Exposure Is Visualised by Stimulated Raman Scattering Microscopy. Int J Mol Sci 18: 894

Busk PK (2014) A tool for design of primers for microRNA-specific quantitative RT-qPCR. BMC Bioinformatics 15: 1–9

Cai D & Liu T (2012) INTRODUCTION: Brain inflammation in metabolic syndrome. Aging (Albany NY) 4: 98–115

Cai R, Sun Y, Qimuge N, Wang G, Wang Y, Chu G, Yu T, Yang G & Pang W (2018) Adiponectin AS lncRNA inhibits adipogenesis by transferring from nucleus to cytoplasm and attenuating Adiponectin mRNA translation. Biochim Biophys Acta - Mol Cell Biol Lipids 1863: 420–432

Catrysse L & van Loo G (2017) Inflammation and the Metabolic Syndrome: The Tissue-Specific Functions of NF-κB. Trends Cell Biol 27: 417–429

Craske MG, Stein MB, Eley TC, Milad MR, Holmes A, Rapee RM & Wittchen H-U (2017) Anxiety disorders. Nat Rev Dis Prim 3: 1–18

Cui X, Niu W, Kong L, He M, Jiang K, Chen S, Zhong A, Zhang Q, Li W, Lu J, et al (2017) Long noncoding RNA as an indicator differentiating schizophrenia from major depressive disorder and generalized anxiety disorder in nonpsychiatric hospital. Biomark Med 11: 221–228

Deniz E & Erman B (2017) Long noncoding RNA (lincRNA), a new paradigm in gene expression control. Funct Integr Genomics 17: 135–143

Doepker C, Lieberman HR, Smith AP, Peck JD, El-Sohemy A & Welsh BT (2016) Caffeine: Friend or Foe? Annu Rev Food Sci Technol 7: 117–137

Dornellas APS, Boldarine VT, Pedroso AP, Carvalho LOT, de Andrade IS, Vulcani-Freitas TM, dos Santos CCC, do Nascimento CM d. PO, Oyama LM & Ribeiro EB (2018) High-fat feeding improves anxiety-type behavior induced by ovariectomy in rats. Front Neurosci 12: 1–13

Dulloo AG (2011) The search for compounds that stimulate thermogenesis in obesity management: From pharmaceuticals to functional food ingredients. Obes Rev 12: 866–883

Dutheil S, Ota KT, Wohleb ES, Rasmussen K & Duman RS (2016) High-Fat Diet Induced Anxiety and Anhedonia: Impact on Brain Homeostasis and Inflammation. Neuropsychopharmacology 41: 1874–1887

Egan RJ, Bergner CL, Hart PC, Cachat JM, Canavello PR, Elegante MF, Elkhayat SI, Bartels BK, Tien AK, Tien DH, et al (2009) Understanding behavioral and physiological phenotypes of stress and anxiety in zebrafish. Behav Brain Res 205: 38–44

Elhaik Goldman S, Goez D, Last D, Naor S, Liraz Zaltsman S, Sharvit-Ginon I, Atrakchi-Baranes D, Shemesh C, Twitto-Greenberg R, Tsach S, et al (2018) High-fat diet protects the blood-brain barrier in an Alzheimer’s disease mouse model. Aging Cell 17: e12818

Fan K qi, Li Y yuan, Wang H li, Mao X tao, Guo J xin, Wang F, Huang L jie, Li Y ning, Ma X yu, Gao Z jun, et al (2019) Stress-Induced Metabolic Disorder in Peripheral CD4+ T Cells Leads to Anxiety-like Behavior. Cell 179: 864–879.e19

Felger JC (2017) Imaging the role of inflammation in mood and anxiety-related disorders. Curr Neuropharmacol 15: 533–558

Filetti V, La Ferlita A, Di Maria A, Cardile V, Graziano ACE, Rapisarda V, Ledda C, Pulvirenti A & Loreto C (2022) Dysregulation of microRNAs and tRNA-derived ncRNAs in mesothelial and mesothelioma cell lines after asbestiform fiber exposure. Sci Rep 12: 1–21

Filiano AJ, Xu Y, Tustison NJ, Marsh RL, Baker W, Smirnov I, Overall CC, Gadani SP, Turner SD, Weng Z, et al (2016) Unexpected role of interferon-γ 3 in regulating neuronal connectivity and social behaviour. Nature 535: 425–429

Finger BC, Dinan TG & Cryan JF (2011) High-fat diet selectively protects against the effects of chronic social stress in the mouse. Neuroscience 192: 351–360

Fontana BD, Cleal M & Parker MO (2020) Female adult zebrafish (Danio rerio) show higher levels of anxiety-like behavior than males, but do not differ in learning and memory capacity. Eur J Neurosci 52: 2604–2613

Fontana BD, Mezzomo NJ, Kalueff A V. & Rosemberg DB (2018) The developing utility of zebrafish models of neurological and neuropsychiatric disorders: A critical review. Exp Neurol 299: 157– 171

Fourrier C, Bosch-Bouju C, Boursereau R, Sauvant J, Aubert A, Capuron L, Ferreira G, Layé S & Castanon N (2019) Brain tumor necrosis factor-α mediates anxiety-like behavior in a mouse model of severe obesity. Brain Behav Immun 77: 25–36

Francés DE, Motiño O, Agrá N, González-Rodríguez Á, Fernández-Álvarez A, Cucarella C, Mayoral R, Castro-Sánchez L, García-Casarrubios E, Boscá L, et al (2015) Hepatic cyclooxygenase-2 expression protects against diet-induced steatosis, obesity, and insulin resistance. Diabetes 64: 1522–1531

Friedrich RW, Jacobson GA & Zhu P (2010) Circuit Neuroscience in Zebrafish. Curr Biol 20: R371–R381

Fu SP, Hong H, Lu SF, Hu CJ, Xu HX, Li Q, Yu ML, Ou C, Meng JZ, Wang TL, et al (2017) Genome-wide regulation of electro-acupuncture on the neural Stat5-loss-induced obese mice. PLoS One 12: 1–19

Gainey SJ, Kwakwa KA, Bray JK, Pillote MM, Tir VL, Towers AE & Freund GG (2016) Short-term high-fat diet (HFD) induced anxiety-like behaviors and cognitive impairment are improved with treatment by glyburide. Front Behav Neurosci 10: 1–12

Gautier Y, Bergeat D, Serrand Y, Réthoré N, Mahérault M, Malbert CH, Meurice P, Coquery N, Moirand R & Val-Laillet D (2020) Western diet, obesity and bariatric surgery sequentially modulated anxiety, eating patterns and brain responses to sucrose in adult Yucatan minipigs. Sci Rep 10: 1–18

Hanin G, Yayon N, Tzur Y, Haviv R, Bennett ER, Udi S, Krishnamoorthy YR, Kotsiliti E, Zangen R, Efron B, et al (2018) miRNA-132 induces hepatic steatosis and hyperlipidaemia by synergistic multitarget suppression. Gut 67: 1124–1134

Hiles SA, Révész D, Lamers F, Giltay E & Penninx BWJH (2016) Bidirectional Prospective Associations of Metabolic Syndrome Components With Depression, Anxiety, and Antidepressant Use. Depress Anxiety 33: 754–764

Hogue CW, Stearns JD, Colantuoni E, Robinson KA, Stierer T, Mitter N, Pronovost PJ & Needham DM (2009) The impact of obesity on outcomes after critical illness: A meta-analysis. Intensive Care Med 35: 1152–1170

Howe K, Clark MD, Torroja CF, Torrance J, Berthelot C, Muffato M, Collins JE, Humphray S, McLaren K, Matthews L, et al (2013) The zebrafish reference genome sequence and its relationship to the human genome. Nature 496: 498–503

Huang PL (2009) A comprehensive definition for metabolic syndrome. Dis Model Mech 2: 231–237

Hutagalung R, Marques J, Kobylka K, Zeidan M, Kabisch B, Brunkhorst F, Reinhart K & Sakr Y (2011) The obesity paradox in surgical intensive care unit patients. Intensive Care Med 37: 1793–1799

Iacomino G & Siani A (2017) Role of microRNAs in obesity and obesity-related diseases. Genes Nutr 12: 1–16

Issler O & Chen A (2015) Determining the role of microRNAs in psychiatric disorders. Nat Rev Neurosci 16: 201–212

Ke W, Reed JN, Yang C, Higgason N, Rayyan L, Wahlby C, Carpenter AE, Civelek M & O’Rourke EJ (2021) Genes in human obesity loci are causal obesity genes in C. elegans

Khyzha N, Khor M, DiStefano P V., Wang L, Matic L, Hedin U, Wilson MD, Maegdefessel L & Fish JE (2019) Regulation of CCL2 expression in human vascular endothelial cells by a neighboring divergently transcribed long noncoding RNA. Proc Natl Acad Sci U S A 116: 16410–16419

Kim HK, Yeom JH & Kay MA (2020) Transfer RNA-Derived Small RNAs: Another Layer of Gene Regulation and Novel Targets for Disease Therapeutics. Mol Ther 28: 2340–2357

Kolshus E, Dalton VS, Ryan KM & McLoughlin DM (2014) When less is more - microRNAs and psychiatric disorders. Acta Psychiatr Scand doi:10.1111/acps.12191 [PREPRINT]

Lenze EJ, Mantella RC, Shi P, Goate AM, Nowotny P, Butters MA, Andreescu C, Thompson PA & Rollman BL (2011) Elevated cortisol in older adults with generalized anxiety disorder is reduced by treatment: A placebo-controlled evaluation of escitalopram. Am J Geriatr Psychiatry 19: 482–490

Lieschke GJ & Currie PD (2007) Animal models of human disease: Zebrafish swim into view. Nat Rev Genet 8: 353–367

Lo KA, Huang S, Walet C, Zhang Z, Leow M, Meihui L & Sun L (2018) Adipocyte long noncoding RNA transcriptome analysis of obese mice identified Lnc-leptin which regulates Leptin. Diabetes: 1– 51

Loher P, Telonis AG & Rigoutsos I (2017) MINTmap: Fast and exhaustive profiling of nuclear and mitochondrial tRNA fragments from short RNA-seq data. Sci Rep 7: 1–20

López Nadal A, Ikeda-Ohtsubo W, Sipkema D, Peggs D, McGurk C, Forlenza M, Wiegertjes GF & Brugman S (2020) Feed, Microbiota, and Gut Immunity: Using the Zebrafish Model to Understand Fish Health. Front Immunol 11

Luyer MD, Greve JWM, Hadfoune M, Jacobs JA, Dejong CH & Buurman WA (2005) Nutritional stimulation of cholecystokinin receptors inhibits inflammation via the vagus nerve. J Exp Med 202: 1023–1029

Madrer N & Soreq H (2020) Cholino-ncRNAs modulate sex-specific- and age-related acetylcholine signals. FEBS Lett 594: 2185–2198

Marín-Juez R, Jong-Raadsen S, Yang S & Spaink HP (2014) Hyperinsulinemia induces insulin resistance and immune suppression via Ptpn6/Shp1 in zebrafish. J Endocrinol 222: 229–241

Mathew BA, Katta M, Ludhiadch A, Singh P & Munshi A (2022) Role of tRNA-Derived Fragments in Neurological Disorders: a Review. Mol Neurobiol

McNabb HJ, Zhang Q & Sjögren B (2020) Emerging Roles for Regulator of G Protein Signaling 2 in (Patho)physiology. Mol Pharmacol 98: 751–760

McNeilly AD, Stewart CA, Sutherland C & Balfour DJK (2015) High fat feeding is associated with stimulation of the hypothalamic-pituitary-adrenal axis and reduced anxiety in the rat. Psychoneuroendocrinology 52: 272–280

Di Meco A, Jelinek J, Lauretti E, Curtis ME, Issa JPJ & Praticό D (2019) Gestational high fat diet protects 3xTg offspring from memory impairments, synaptic dysfunction, and brain pathology. Mol Psychiatry: 7006–7019

Di Meco A & Praticò D (2019) Early-life exposure to high-fat diet influences brain health in aging mice. Aging Cell 18: 1–9

Meguro S, Hosoi S & Hasumura T (2019) High-fat diet impairs cognitive function of zebrafish. Sci Rep 9: 1–9

Meier SM & Deckert J (2019) Genetics of Anxiety Disorders. Curr Psychiatry Rep 21

Mendes NF, Castro G, Guadagnini D, Tobar N, Cognuck SQ, Elias LLK, Boer PA & Prada PO (2017) Knocking down amygdalar PTP1B in diet-induced obese rats improves insulin signaling/action, decreases adiposity and may alter anxiety behavior. Metabolism 70: 1–11

Meydan C, Shenhar-Tsarfaty S & Soreq H (2016) MicroRNA Regulators of Anxiety and Metabolic Disorders. Trends Mol Med 22: 798–812

Michopoulos V, Powers A, Gillespie CF, Ressler KJ & Jovanovic T (2017) Inflammation in Fear-and Anxiety-Based Disorders: PTSD, GAD, and beyond. Neuropsychopharmacology 42: 254–270

Minchin JEN & Rawls JF (2011) In vivo Analysis of White Adipose Tissue in Zebrafish. Methods Cell Biol 105: 63–86

Misri S & Kendrick K (2007) Treatment of Perinatal Mood and Anxiety Disorders: A Review. Can J Psychiatry 52: 489–498

Mraz M & Haluzik M (2014) The role of adipose tissue immune cells in obesity and low-grade inflammation. J Endocrinol 222: 113–127

Murphy CP & Singewald N (2019) Role of MicroRNAs in Anxiety and Anxiety-Related Disorders. In Brain Imaging in Behavioral Neuroscience pp 289–320.

Nguyen M, Yang E, Neelkantan N, Mikhaylova A, Arnold R, Poudel MK, Stewart AM & Kalueff A V. (2013) Developing ‘integrative’ zebrafish models of behavioral and metabolic disorders. Behav Brain Res 256: 172–187

Nousen EK, Franco JG & Sullivan EL (2014) Unraveling the mechanisms responsible for the comorbidity between metabolic syndrome and mental health disorders. Neuroendocrinology 98: 254–266

Novoa B & Figueras A (2012) Zebra fish Model for studying Innate Immunity. In Current Topics in Innate Immunity II pp 253–275.

Ogrodnik M, Zhu Y, Langhi LGP, Tchkonia T, Krüger P, Fielder E, Victorelli S, Ruswhandi RA, Giorgadze N, Pirtskhalava T, et al (2019) Obesity-Induced Cellular Senescence Drives Anxiety and Impairs Neurogenesis. Cell Metab: 1–17

Picolo VL, Quadros VA, Canzian J, Grisolia CK, Goulart JT, Pantoja C, de Bem AF & Rosemberg DB (2021) Short-term high-fat diet induces cognitive decline, aggression, and anxiety-like behavior in adult zebrafish. Prog Neuro-Psychopharmacology Biol Psychiatry 110: 110288

Pierce GL, Kalil GZ, Ajibewa T, Holwerda SW, Persons J, Moser DJ & Fiedorowicz JG (2017) Anxiety independently contributes to elevated inflammation in humans with obesity. Obesity 25: 286– 289

Poitou C, Perret C, Mathieu F, Truong V, Blum Y, Durand H, Alili R, Chelghoum N, Pelloux V, Aron-Wisnewsky J, et al (2015) Bariatric surgery induces disruption in inflammatory signaling pathways mediated by immune cells in adipose tissue: A RNA-seq study. PLoS One 10: 1–23

Pontzer H, Yamada Y, Sagayama H, Ainslie PN, Andersen LF, Anderson LJ, Arab L, Baddou I, Bedu-Addo K, Blaak EE, et al (2021) Daily energy expenditure through the human life course. Science (80-) 373: 808–812

Pritchard CC, Cheng HH & Tewari M (2012) MicroRNA profiling: approaches and considerations. Nat Rev Genet 13: 358–369

Provenzano F & Deleidi M (2021) Reassessing neurodegenerative disease: immune protection pathways and antagonistic pleiotropy. Trends Neurosci 44: 771–780

Van Reedt Dortland AKB, Giltay EJ, Van Veen T, Zitman FG & Penninx BWJH (2013) Longitudinal relationship of depressive and anxiety symptoms with dyslipidemia and abdominal obesity. Psychosom Med 75: 83–89

Richendrfer H, Pelkowski SD, Colwill RM & Creton R (2012) On the edge: Pharmacological evidence for anxiety-related behavior in zebrafish larvae. Behav Brain Res 228: 99–106

Robinson MD, McCarthy DJ & Smyth GK (2009) edgeR: A Bioconductor package for differential expression analysis of digital gene expression data. Bioinformatics 26: 139–140

Robinson OJ, Pike AC, Cornwell B & Grillon C (2019) The translational neural circuitry of anxiety. J Neurol Neurosurg Psychiatry 90: 1353–1360

Rodríguez-Calvo R, Tajes M & Vázquez-Carrera M (2017) The NR4A subfamily of nuclear receptors: potential new therapeutic targets for the treatment of inflammatory diseases. Expert Opin Ther Targets 21: 291–304

Rohm TV., Meier DT, Olefsky JM & Donath MY (2022) Inflammation in obesity, diabetes, and related disorders. Immunity 55: 31–55

Rosa LV, Ardais AP, Costa FV, Fontana BD, Quadros VA, Porciúncula LO & Rosemberg DB (2018) Different effects of caffeine on behavioral neurophenotypes of two zebrafish populations. Pharmacol Biochem Behav 165: 1–8

Rosace D, López J & Blanco S (2020) Emerging roles of novel small non-coding regulatory RNAs in immunity and cancer. RNA Biol 17: 1196–1213

Ruiz-Oliveira J, Silva PF & Luchiari AC (2019) Coffee time: Low caffeine dose promotes attention and focus in zebrafish. Learn Behav 47: 227–233

Serikuly N, Alpyshov ET, Wang D, Wang J, Yang L, Hu G, Yan D, Demin KA, Kolesnikova TO, Galstyan D, et al (2020) Effects of acute and chronic arecoline in adult zebrafish: Anxiolytic-like activity, elevated brain monoamines and the potential role of microglia. Prog Neuro-Psychopharmacology Biol Psychiatry: 109977

Shimada-Sugimoto M, Otowa T & Hettema JM (2015) Genetics of anxiety disorders: Genetic epidemiological and molecular studies in humans. Psychiatry Clin Neurosci 69: 388–401

Singleman C & Holtzman NG (2014) Growth and Maturation in the Zebrafish, *Danio Rerio* : A Staging Tool for Teaching and Research. Zebrafish 11: 396–406

Soreq H (2015) Checks and balances on cholinergic signaling in brain and body function. Trends Neurosci 38: 448–458

Soto M, Herzog C, Pacheco JA, Fujisaka S, Bullock K, Clish CB & Kahn CR (2018) Gut microbiota modulate neurobehavior through changes in brain insulin sensitivity and metabolism. Mol Psychiatry 23: 2287–2301

Spadaro PA, Flavell CR, Widagdo J, Ratnu VS, Troup M, Ragan C, Mattick JS & Bredy TW (2015) Long Noncoding RNA-Directed Epigenetic Regulation of Gene Expression Is Associated With Anxiety-like Behavior in Mice. Biol Psychiatry 78: 848–859

Sudre CH, Murray B, Varsavsky T, Graham MS, Penfold RS, Bowyer RC, Pujol JC, Klaser K, Antonelli M, Canas LS, et al (2021) Attributes and predictors of long COVID. Nat Med 27: 626–631

Tang F, Wang G & Lian Y (2017) Association between anxiety and metabolic syndrome: A systematic review and meta-analysis of epidemiological studies. Psychoneuroendocrinology 77: 112–121

Tang R, Dodd A, Lai D, McNabb WC & Love DR (2007) Validation of zebrafish (Danio rerio) reference genes for quantitative real-time RT-PCR normalization. Acta Biochim Biophys Sin (Shanghai) 39: 384–390

Tran S, Fulcher N, Nowicki M, Desai P, Tsang B, Facciol A, Chow H & Gerlai R (2017) Time-dependent interacting effects of caffeine, diazepam, and ethanol on zebrafish behaviour. Prog Neuro-Psychopharmacology Biol Psychiatry 75: 16–27

Türkoğlu M, Baran A, Sulukan E, Ghosigharehagaji A, Yildirim S, Ceyhun HA, Bolat İ, Arslan M & Ceyhun SB (2022) The potential effect mechanism of high-fat and high-carbohydrate diet-induced obesity on anxiety and offspring of zebrafish. Eat Weight Disord - Stud Anorexia, Bulim Obes 27: 163–177

Walsh K, Furey WJ & Malhi N (2021) Narrative review: COVID-19 and pediatric anxiety. J Psychiatr Res 144: 421–426

Wang X, Yang Y, Tan X, Mao X, Wei D, Yao Y, Jiang P, Mo D, Wang T & Yan F (2019) Identification of tRNA-Derived Fragments Expression Profile in Breast Cancer Tissues. Curr Genomics 20: 199–213

Welty FK, Alfaddagh A & Elajami TK (2016) Targeting inflammation in metabolic syndrome. Transl Res 167: 257–280

Westerfield M (2007) THE ZEBRAFISH BOOK, 5th Edition; A guide for the laboratory use of zebrafish (Danio rerio) 5th ed. University of Oregon Press

Wickham H (2016) ggplot2 Cham: Springer International Publishing

Williams GC (1957) Pleiotropy, Natural Selection, and the Evolution of Senescence. Evolution (N Y) 11: 398–411

Wilson MR, Petrie JE, Shaw MW, Hu C, Oakley CM, Woods SJ, Patel B V., O’Dea KP & Takata M (2017) High-fat feeding protects mice from ventilator-induced lung injury, via neutrophil-independent mechanisms. Crit Care Med 45: e831–e839

Winek K, Lobentanzer S, Nadorp B, Dubnov S, Dames C, Jagdmann S, Moshitzky G, Hotter B, Meisel C, Greenberg DS, et al (2020) Transfer RNA fragments replace microRNA regulators of the cholinergic poststroke immune blockade. Proc Natl Acad Sci U S A 117: 32606–32616

Wong K, Elegante M, Bartels B, Elkhayat S, Tien D, Roy S, Goodspeed J, Suciu C, Tan J, Grimes C, et al (2010) Analyzing habituation responses to novelty in zebrafish (Danio rerio). Behav Brain Res 208: 450–457

Wong RY & Godwin J (2015) Neurotranscriptome profiles of multiple zebrafish strains. Genomics Data 5: 206–209

Wong RY, Perrin F, Oxendine SE, Kezios ZD, Sawyer S, Zhou L, Dereje S & Godwin J (2012) Comparing behavioral responses across multiple assays of stress and anxiety in zebrafish (Danio rerio). Behaviour 149: 1205–1240

Xia G, Han Y, Meng F, He Y, Srisai D, Farias M, Dang M, Palmiter RD, Xu Y & Wu Q (2021) Reciprocal control of obesity and anxiety–depressive disorder via a GABA and serotonin neural circuit. Mol Psychiatry 26: 2837–2853

Yeh C-M, Glöck M & Ryu S (2013) An Optimized Whole-Body Cortisol Quantification Method for Assessing Stress Levels in Larval Zebrafish. PLoS One 8: e79406

Yin J, Zhang J & Lu Q (2017) The role of basic leucine zipper transcription factor E4BP4 in the immune system and immune-mediated diseases. Clin Immunol 180: 5–10

Yoon G, Cho KA, Song J & Kim YK (2019) Transcriptomic analysis of high fat diet fed mouse brain cortex. Front Genet 10: 1–13

Zang L, Maddison LA & Chen W (2018) Zebrafish as a Model for Obesity and Diabetes. Front Cell Dev Biol 6: 1–13

Zang L, Shimada Y, Nakayama H, Kim Y, Chu DC, Juneja LR, Kuroyanagi J & Nishimura N (2019) RNA-seq based transcriptome analysis of the anti-obesity effect of green tea extract using zebrafish obesity models. Molecules 24: 1–11

Zheng X, Dai W, Chen X, Wang K, Zhang W, Liu L & Hou J (2015) Caffeine reduces hepatic lipid accumulation through regulation of lipogenesis and ER stress in zebrafish larvae. J Biomed Sci 22: 1–12

Zhu C, Hui L, Zheng K, Liu L, Liu J & Lv W (2020) Silencing of RGS2 enhances hippocampal neuron regeneration and rescues depression-like behavioral impairments through activation of cAMP pathway. Brain Res 1746: 147018

Zorbaz T, Madrer N & Soreq H (2022) Cholinergic blockade of neuroinflammation: From tissue to RNA regulators. Neuronal Signal 6: 1–15

